# HCN channels enhance synchrony propagation in heterogeneous synfire chains

**DOI:** 10.64898/2026.05.28.728444

**Authors:** Sarang Saini, Rishikesh Narayanan

**Affiliations:** Cellular Neurophysiology Laboratory, Molecular Biophysics Unit, Indian Institute of Science, Bangalore, India

**Keywords:** computational model, degeneracy, HCN channel, neocortex, network-to-network variability, neural-circuit heterogeneities, synchrony propagation

## Abstract

**Motivation:** Information flow and temporal coding in cortical circuits depend critically on the reliable transmission of precisely timed synchronous spike patterns. Although cortical assemblies achieve such transmission despite pronounced intrinsic heterogeneities and stochastic high-conductance states, the mechanisms underlying effective synchrony propagation under *in vivo* conditions remain poorly understood.

**Methodology:** In this study, we address this gap using large-scale, conductance-based models of excitatory and inhibitory neurons organized into feedforward synfire chains operating in noisy, high-conductance regimes. Using independent stochastic search algorithms, we first identified physiologically valid heterogeneous populations of cortical neurons. Both excitatory and inhibitory populations exhibited cellular-scale degeneracy, whereby distinct combinations of biophysically identified molecular components produced signature physiological characteristics. We then constructed synfire chains with varying degrees of heterogeneity using these populations and assessed the propagation of different spike packets across neuronal assemblies.

**Results:** We found synchrony propagation to be inherently probabilistic, revealing a stochastic separatrix that separated input patterns that consistently succeeded from those that consistently failed in propagation. The stochastic nature of this separatrix highlighted a critical role for background synaptic fluctuations, defining a regime in which identical inputs alternately propagated or failed across trials solely due to stochastic background activity. Comparing networks with different degrees of intrinsic heterogeneity, we found that increasing heterogeneity did not alter mean propagation efficacy but reduced network-to-network variability, indicating a stabilizing role for intrinsic diversity. Strikingly, when we tested the impact of neuronal intrinsic properties on synchrony propagation, hyperpolarization-activated cyclic nucleotide-gated (HCN) channels emerged as robust enhancers of synchrony propagation across all heterogeneity regimes. Mechanistically, the slow restorative kinetics of HCN conductances narrowed the temporal window for spike initiation, sharpening output synchrony, and improving propagation reliability. This effect was abolished when HCN kinetics were accelerated, underscoring the importance of the slow negative feedback mediated by these channels.

**Implications:** Together, our analyses identify HCN channels as key regulators of synchronous information transfer and reveal strong interactions among intrinsic conductances, input characteristics, neuronal heterogeneity, and stochastic background activity in shaping cortical synchrony propagation. The ability of diverse cellular and network configurations to achieve similar propagation efficacy further highlights degeneracy as a fundamental principle governing robust and flexible neural computation.

## INTRODUCTION

Information flow through cortical circuits depends critically on their ability to generate, transform, and reliably transmit precisely timed patterns of action potentials, despite widespread heterogeneities in cellular and network components and in the presence of pervasive background noise. Temporal codes, where information is embedded in the relative timing and synchrony of spikes, play a central role in sensory processing, navigation, cognition, and memory (O’Keefe and Recce, 1993; Quian Quiroga and Panzeri, 2009; Buzsaki, 2010; Ratte et al., 2013; Panzeri et al., 2017; Xie et al., 2024). For such timing-based representations to be functionally meaningful, neural circuits must not only support the generation of synchronous spike patterns but also sustain stable propagation of synchrony across successive stages of processing. Feedforward propagation across neural assemblies in the *temporal dimension* is represented by synfire chains, a theoretical framework that has been useful in understanding how synchrony propagates across cortical structures (Abeles, 1982, 2009). Prior computational and experimental studies have demonstrated that synchronous spike packets can propagate through such networks, with propagation governed by the size and temporal dispersion of input spike volleys. These studies have led to the concept of a separatrix that divides propagating from non-propagating input regimes and have demonstrated how synaptic connectivity and input structure influence synchrony propagation (Diesmann et al., 1999; Bartos et al., 2001; Gewaltig et al., 2001; van Rossum et al., 2002; Reyes, 2003; Ikegaya et al., 2004; Kumar et al., 2008, 2010; Jahnke et al., 2013; Moldakarimov et al., 2015; Ashhad and Feldman, 2020; Ashhad et al., 2023).

However, most work on cortical synchrony propagation has relied on simplified neuron models, homogeneous assembly architectures, or background synaptic inputs that do not account for high-conductance states. Specifically, as prior analyses have predominantly focused on the characteristics of input packets, the specific roles of intrinsic properties of cortical neurons on synchrony propagation have not been studied thoroughly. In doing this, it is essential that single-neuron models account for the characteristic intrinsic properties of excitatory and inhibitory neurons through biophysical modeling of the several ion channels that shape these properties. In the absence of conductance-based models for neurons, the impact of intrinsic characteristics or specific neuronal ion channels on synchrony propagation cannot be studied (Hong et al., 2012; Ratte et al., 2013). Second, within-type heterogeneity is a defining feature of cortical circuits (Shamir and Sompolinsky, 2006; Padmanabhan and Urban, 2010; Hay et al., 2011; Tripathy et al., 2013; Cembrowski and Spruston, 2019; Tzilivaki et al., 2019; Moradi Chameh et al., 2021; Rich et al., 2022; Planert et al., 2025; Dahmen et al., 2026). Individual excitatory and inhibitory neurons within the same subtype differ widely in their intrinsic excitability, ion-channel expression, and firing dynamics. At the network level, this cellular diversity translates into pronounced heterogeneity across circuit instantiations, raising a fundamental question on how robust synchrony propagation is to intrinsic neuronal heterogeneity. Third, cortical neurons *in vivo* exhibit signature high-conductance states as a consequence of stochastic background synaptic inputs that open excitatory and inhibitory receptors. High-conductance states are characterized by synaptic noise and significant reduction in neuronal gain (Chance et al., 2002; Destexhe et al., 2003; Mishra and Narayanan, 2015), and thus could alter spiking dynamics and synchrony propagation in cortical structures. Together, synchrony propagation in biologically realistic conductance-based heterogeneous neural assemblies that operate in stochastic high-conductance states has remained unexplored.

To address this question, here, we combine large-scale conductance-based modeling of single neurons with a population-of-networks approach to assess synchrony propagation in stochastic heterogeneous synfire chains. We construct physiologically validated heterogeneous populations of cortical excitatory and inhibitory neurons using multi-parametric, multi-objective stochastic search, ensuring that these biophysically grounded conductance-based model neurons capture experimentally observed heterogeneities in their physiological characteristics. We assembled these neurons into feedforward networks of excitatory-inhibitory assemblies operating in balanced high-conductance states, allowing us to systematically probe how synchronous spike packets propagate across noisy, heterogeneous cortical circuits.

Our analyses fundamentally demonstrated that cortical circuits can robustly sustain precise temporal coding and efficacious synchrony propagation despite intrinsic heterogeneities and stochastic high-conductance states, revealing three central findings. First, synchrony propagation in high-conductance networks is inherently probabilistic, giving rise to a stochastic separatrix: a zone of uncertainty where identical inputs may succeed or fail across trials owing solely to fluctuations in background activity. Second, increasing the degree of intrinsic heterogeneity does not impair average propagation efficacy but reduces network-to-network variability, highlighting a stabilizing role for cellular diversity in synchrony propagation. Third, and most strikingly, we find that HCN channels robustly and consistently enhance synchrony propagation across network instantiations, independent of the degree of heterogeneity. Mechanistically, we find this enhancement to arise from the slow restorative dynamics of HCN channels, which sharpen output spike timing by narrowing the window for spike generation.

Together, our results identify HCN channels as active regulators of synchrony propagation and uncover strong interactions between afferent input characteristics, intrinsic conductances, neuronal heterogeneity, and background noise in shaping synchronous information flow. By demonstrating that similar propagation efficacy can emerge from disparate cellular and network configurations, our findings point to degeneracy as a fundamental organizing principle underlying robust synchrony transmission in cortical circuits. Our analyses also suggest that neuromodulatory and activity-dependent control of intrinsic conductances provides a powerful and flexible mechanism for dynamically gating synchronous activity across the cortex.

## METHODS

Our goal in this study was to understand how synchrony propagates across cortical feedforward networks and how this propagation depends on intrinsic neuronal properties and heterogeneities therein. The overall experimental design towards achieving this was as follows. We first generated independent heterogeneous populations of cortical excitatory and inhibitory neurons. Excitatory neurons were modeled based on the characteristics of layer 5 cortical pyramidal neurons, which form a principal excitatory neuronal population of cortical columns. Inhibitory neurons were modeled based on cortical basket cells, the representative fast-spiking inhibitory interneurons that strongly influence cortical activity. We modeled both neuronal subtypes as single-compartment cylindrical structures endowed with a diverse set of voltage-gated and calcium-dependent ion channels expressed by these neuronal subtypes (Fig. 1*A*). The reduced-model formulation was essential to keep the computational complexity in check, given that each of the several networks was built with 1250 conductance-based neuronal models (Fig. 1*C*), each tested across multiple trials and several input patterns.

**Figure 1:**
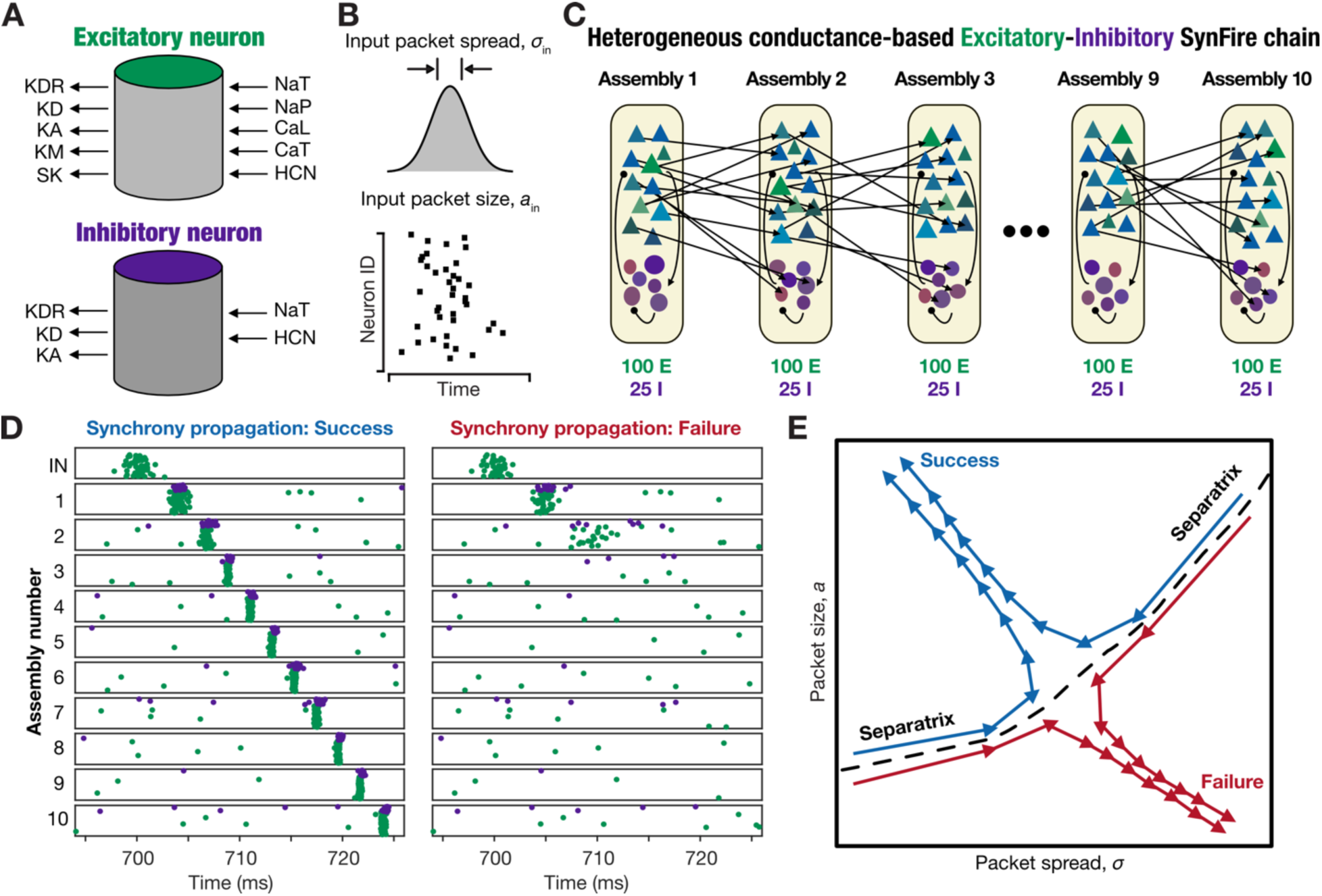
Illustrations of single-neuron models, feedforward network architecture, propagation success, and separatrix dividing propagation success and failure. (A) Schematic showing the ion channels incorporated into excitatory (top) and inhibitory (bottom) neuronal models. All inward currents are to the right and represented with an inward arrow and all outward currents are to the left and represented with an outward arrow (B) Input received by the network. The input spike packet characterized by its size (*a*_*in*_) and spread (*σ*_*in*_). (C) Schematic of the feedforward synfire network, where each of 10 assemblies contained a population of 100 excitatory and 25 inhibitory neurons. (D) Example raster plots across the 10 assemblies showing propagating (left) and non-propagating (right) input volleys. (E) Schematic state-space representation illustrating the separatrix that separates propagating from non-propagating input packets. Success implies that the spike packet size in the last assembly is large with low spread (top left of the state-space), whereas failure depicts a scenario where the packet size there is small with high spread (bottom right of the state-space). Input volleys with relatively smaller packet sizes need low spread for successful propagation across all assemblies (each arrow represents a traversal across two successive assemblies), while those with relatively larger packet sizes can successfully propagate even with relatively high spread. The gradual increase in the value of *a* with increasing *σ* may be noticed along the separatrix (Diesmann et al., 1999).

To match within-subtype heterogeneities in our excitatory and inhibitory neuronal model populations with intrinsic variability observed in respective electrophysiological recordings, we implemented two independent multi-parametric multi-objective stochastic search (MPMOSS) algorithms, one for each neuronal subtype. The model neurons were used to construct populations of feedforward networks with different degrees of heterogeneities, with networks comprising 10 assemblies (Fig. 1*C*). Each assembly contained 100 excitatory and 25 inhibitory neurons that projected to the next assembly in a feedforward sequence, allowing us to examine how synchronous spike packets with different characteristics propagated across successive assemblies. To replicate *in vivo*-like background synaptic activity, all neurons in the network received thousands of stochastic excitatory and inhibitory synaptic inputs that yielded synaptic noise and high-conductance state. Using this experimental design, we examined how synchrony propagation is shaped by characteristics of the input pattern, neuronal intrinsic properties, and different degrees of within-type intrinsic heterogeneity. In assessing the impact of neuronal intrinsic properties, we systematically scaled HCN-channel density across the neuronal population and compared propagation across networks with different densities of HCN channels and several degrees of neuronal heterogeneity. The population-of-networks approach involving heterogeneous populations of each neuronal subtype allowed us to systematically study how intrinsic conductances and variability in neuronal excitability influenced the fidelity of synchrony propagation in cortical feedforward circuits. In what follows, we elaborate details associated with this overall experimental design.

### Biophysical composition and electrophysiological measurements of excitatory neurons

Excitatory cortical neurons were modeled as single-compartment cylindrical structures with 70 µm diameter and 70 µm length (Fig. 1*A*). Each neuron was endowed with 11 ion-channel conductances whose kinetics were adopted from a previous computational model (Hay et al., 2011): leak (represented by *R*_*m*_ = 1/*g*_*leak*_), transient sodium (NaT), persistent sodium (NaP), delayed-rectifier potassium (KDR), *D*-type potassium (KD), *M*-type potassium (KM), *A*-type potassium (KA), *L*-type calcium (CaL), *T*-type calcium (CaT), hyperpolarization-activated cyclic nucleotide-gated (HCN), and small-conductance calcium-activated potassium (SK) channel conductances (Fig. 1*A*; Table 1). All ion-channel gating kinetics were based on the Hodgkin-Huxley (Hodgkin and Huxley, 1952) formalism. Whereas sodium, potassium, and HCN currents were modeled with the Ohmic formulation, calcium currents used the Goldman-Hodgkin-Katz (GHK) formulation (Goldman, 1943; Hodgkin and Katz, 1949) to account for the steep transmembrane concentration gradient in the calcium ion. The reversal potentials for the Na^+^, K^+^, and HCN conductances were set to 50 mV, −90 mV, and −20 mV, respectively. The calcium-handling mechanism, responsible for buffering and pumping cytosolic calcium [*Ca*]_*c*_, was modeled using the following first-order kinetics:

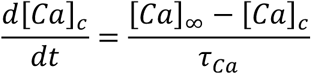

**Table 1:** Bounds on model parameters used for MPMOSS for excitatory neurons.

|  | Model parameter | Lower Bound | Upper Bound |
| --- | --- | --- | --- |
| 1 | Specific membrane resistance, $R_m$ (kΩ cm<sup>2</sup>) | 25 | 45 |
| 2 | Transient sodium channel, $\bar{g}_{NaT}$ (mS/cm<sup>2</sup>) | 90 | 150 |
| 3 | Persistent sodium channel, $\bar{g}_{NaP}$ (μS/cm<sup>2</sup>) | 0.047 | 4.7 |
| 4 | Delayed rectifier potassium channel, $\bar{g}_{KDR}$ (mS/cm<sup>2</sup>) | 10 | 30 |
| 5 | HCN channel, $\bar{g}_{HCN}$ (μS/cm<sup>2</sup>) | 5 | 150 |
| 6 | M-type potassium channel, $\bar{g}_{KM}$ (μS/cm<sup>2</sup>) | 35 | 200 |
| 7 | D-type potassium channel, $\bar{g}_{KD}$ (μS/cm<sup>2</sup>) | 0.1 | 5 |
| 8 | A-type potassium channel, $\bar{g}_{KA}$ (mS/cm<sup>2</sup>) | 0.5 | 2 |
| 9 | L-type activated calcium channel, $\bar{g}_{CaL}$ (mS/cm<sup>2</sup>) | 1 | 4.4 |
| 10 | T-type activated calcium channel, $\bar{g}_{CaT}$ (μS/cm<sup>2</sup>) | 10 | 500 |
| 11 | Small conductance, calcium-activated potassium channel, $\bar{g}_{SK}$ (μS/cm<sup>2</sup>) | 5 | 30 |
| 12 | Time constant of cytosolic calcium decay, $\tau_{Ca}$ (ms) | 80 | 140 |

where 𝜏_*Ca*_ defined the decay time constant of cytosolic calcium and [*Ca*]_∞_ was the steady-state value of cytosolic calcium which was fixed to 100 nM.

We employed a set of standard intrinsic electrophysiological measurements (Narayanan and Johnston, 2007, 2008; Hay et al., 2011; Mishra and Narayanan, 2020; Saini and Narayanan, 2025) to characterize and validate excitatory neuron models, spanning both subthreshold and suprathreshold operational regimes. The resting membrane potential (*V*_RMP_) was measured by allowing each model neuron to settle without injected current for 5 seconds and calculating the mean membrane potential over the following 1 second period. All intrinsic measurements were computed after an initial 5-second stabilization period. Sag ratio (*Sag*) was computed from the model’s voltage response to a hyperpolarizing current injection of −100 pA delivered for 1000 ms. Sag ratio was defined as the ratio between the steady-state membrane deflection and the peak hyperpolarization. Input resistance (*R*_in_) was assessed by injecting a series of current steps ranging from −50 pA to +50 pA in increments of 10 pA, each lasting 1 second. For each step, the steady-state voltage response was recorded, and *R*_in_ was obtained as the slope of a linear fit to the resulting voltage-current (*V* − *I*) graph. We computed rheobase current by injecting 1-second step currents beginning at 0 pA, increasing in increments of 5 pA, until the neuron fired at least one action potential. Rheobase current was defined as the smallest current amplitude that elicited at least one action potential.

Action potential waveforms were characterized using several measurements. For these analyses, the first action potential from a trace with a firing frequency of approximately 9 Hz was used across models. The action potential amplitude was defined as the difference between the spike peak and *V*_RMP_. The spike threshold (*V*_*th*_) was identified as the membrane potential at which the first derivative of the action potential waveform (*dV*/*dt*) exceeded 15 V/s. The action potential half-width (𝑇_APHW_) was measured as the time interval between the upward and downward crossings of the voltage level corresponding to half of the action potential amplitude. The afterhyperpolarization (*V*_AHP_) was defined as the difference between *V*_RMP_and the most negative membrane potential reached between the first and the second action potentials. Adaptation was computed from the inter-spike intervals (ISI) in the firing trace. Specifically, in a trace with *N* spikes, there would be *N* − 1 ISI values. Adaptation was computed from these ISI values as:

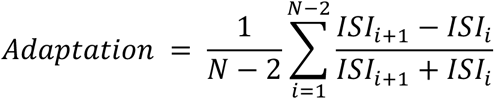

The lower and upper bounds for all these measurements (Table 2) were obtained from previous literature (Schubert et al., 2006; Hay et al., 2011; Herrera et al., 2020; Moradi Chameh et al., 2021).

**Table 2:** Bounds on electrophysiological measurements used for validation of excitatory neurons.

|  | Measurements | Lower Bound | Upper Bound |
| --- | --- | --- | --- |
| 1 | Resting membrane potential, $V_{RMP}$ (mV) | -77 | -65 |
| 2 | Sag ratio, $sag$ | 0.76 | 0.94 |
| 3 | Input resistance, $R_{in}$ (MΩ) | 28 | 204 |
| 4 | Rheobase current, $I_{rheobase}$ (pA) | 55 | 151 |
| 5 | Latency to fire, $T_{1AP}$ (ms) | 21.29 | 65.21 |
| 6 | Peak of the action potential, $V_{AP}$ (mV) | 21 | NA |
| 7 | After hyperpolarization, $V_{AHP}$ (mV) | -64 | -32 |
| 8 | Width of the action potential, $T_{APHW}$ (ms) | 0.8 | 1.8 |
| 9 | Action potential threshold, $V_{th}$ (mV) | -58 | -37 |
| 10 | Firing frequency adaptation, *Adaptation* | -0.0145 | 0.0191 |

We characterized the subthreshold resonance properties of cortical excitatory neurons using impedance analysis. Impedance amplitude (ZAP) profiles were measured using a chirp stimulus consisting of a sinusoidal current of constant amplitude, kept below firing threshold, with frequency linearly spanning 0–15 Hz over 15 s (Chirp15). The impedance as a function of frequency, *Z*(*f*), was computed as the ratio of the Fourier transform of the voltage response to the Fourier transform of the stimulus (Narayanan and Johnston, 2007):

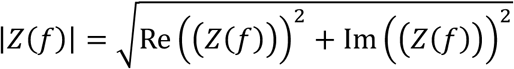

where Re(*Z*(*f*)) and Im(*Z*(*f*)) represent the real and imaginary components of the impedance, respectively. The maximum value of |*Z*(*f*)| across all frequencies was recorded as the maximal impedance amplitude (|*Z*|_*ma*𝑥_) and the frequency at which this maximum occurred was defined as the resonance frequency (*f*_*R*_). Resonance strength (𝑄) was calculated as the ratio of |*Z*|_*ma*𝑥_ to the impedance amplitude at 0.5 Hz.

### Biophysical composition and electrophysiological measurements of inhibitory neurons

Cortical inhibitory neurons were also modeled as single-compartment cylindrical structures with both diameter and length set to 80 µm (Fig. 1*A*). The model incorporated a set of voltage-gated ion channels, whose kinetics and conductance values were adopted from a previous computational modeling study (Tzilivaki et al., 2019). These conductances included a leak channel and 5 voltage-gated ion channels (Fig. 1*A*; Table 3): transient sodium (NaT), delayed-rectifier potassium (KDR), *D*-type potassium (KD), *A*-type potassium (KA), and hyperpolarization-activated cyclic nucleotide-gated (HCN). All conductances were modeled using the Ohmic Hodgkin-Huxley formulation.

**Table 3:** Bounds on model parameters used for MPMOSS for inhibitory neurons.

|  | Model parameter | Lower Bound | Upper Bound |
| --- | --- | --- | --- |
| 1 | Specific membrane resistance, $R_m$ ($\text{k}\Omega \text{ cm}^2$) | 15 | 35 |
| 2 | Transient sodium channel, $\bar{g}_{\text{NaT}}$ ($\text{mS}/\text{cm}^2$) | 10 | 100 |
| 3 | Delayed rectifier potassium channel, $\bar{g}_{\text{KDR}}$ ($\text{mS}/\text{cm}^2$) | 1 | 40 |
| 4 | A-type potassium channel, $\bar{g}_{\text{KA}}$ ($\text{mS}/\text{cm}^2$) | 0.25 | 2.5 |
| 5 | D-type potassium channel, $\bar{g}_{\text{KD}}$ ($\mu\text{S}/\text{cm}^2$) | 10 | 90 |
| 6 | HCN channel, $\bar{g}_{\text{HCN}}$ ($\mu\text{S}/\text{cm}^2$) | 1 | 10 |

A set of intrinsic physiological measurements were computed from interneuron models to characterize and validate their physiology. Resting membrane potential (*V*_RMP_) was measured after allowing the cell to stabilize for 5 s with no current injection and was calculated as the mean membrane potential over a 1-s window between the 5–6 s period. All subsequent intrinsic measurements were computed once the voltage had settled at *V*_RMP_. Sag ratio (*Sag*) was measured as the ratio of the steady-state voltage deflection to the peak voltage deflection in response to a 200 pA hyperpolarizing current step lasting 1000 ms. Action potential firing was quantified by counting the number of spikes evoked by depolarizing current pulses of 50 pA (*f*_50_) and 250 pA (*f*_250_) over 1 s. Input resistance (*R*_in_), a measure of subthreshold excitability, was determined by injecting 1-s current steps ranging from –50 pA to 50 pA in 10 pA increments and recording the corresponding steady-state voltage responses. *R*_in_was calculated as the slope of the resulting voltage-current (*V* − *I*) plot.

Action potential amplitude (*V*_AP_) was defined as the difference between the spike peak and *V*_RMP_. Spike threshold *V*_th_ was identified as the membrane potential at which the first derivative of the action potential waveform (*dV*/*dt*) exceeded 20 V/s. Action potential half-width (𝑇_APHW_) was measured as the duration at which the membrane potential was halfway between *V*_th_ and the spike peak. Afterhyperpolarization (*V*_AHP_) was calculated as the difference between *V*_th_and the most negative membrane potential reached following the spike. Action potential measurements were obtained from the first action potential elicited in response to a 250 pA current pulse injected for a 1-s period. Characteristic ranges of all these measurements (Table 4) were obtained from previous literature (Tzilivaki et al., 2019). We also characterized the subthreshold resonance properties of cortical basket cells, using the same procedure as described for L5 pyramidal neurons, for computing impedance profile (*Z*(*f*)), maximum impedance amplitude (|*Z*|_*ma*𝑥_), resonance frequency (*f*_*R*_) and resonance strength (𝑄).

**Table 4:** Bounds on electrophysiological measurements used for validation of inhibitory neurons.

|  | Measurements | Lower Bound | Upper Bound |
| --- | --- | --- | --- |
| 1 | Resting membrane potential, $V_{\text{RMP}}$ (mV) | -75 | -65 |
| 2 | Standard deviation of resting membrane potential, $\sigma_{\text{RMP}}$ | 0 | 1 |
| 3 | Sag ratio, $Sag$ | 0.90 | 0.98 |
| 4 | Input resistance, $R_{\text{in}}$ ($\text{M}\Omega$) | 70 | 130 |
| 5 | Action potential threshold, $V_{\text{th}}$ (mV) | -41 | -33 |
| 6 | Action potential amplitude, $V_{\text{AP}}$ (mV) | 80 | N/A |
| 7 | Action potential width at half maximum, $T_{\text{APHW}}$ (ms) | N/A | 1 |
| 8 | Afterhyperpolarization, $V_{\text{AHP}}$ (mV) | 20 | N/A |
| 9 | Number of APs for a 50 pA step current for 1000 ms, $f_{50}$ (Hz) | 0 | 0 |
| 10 | Number of APs for a 250 pA step current for 1000 ms, $f_{250}$ (Hz) | 30 | 50 |

### Multi-parametric multi-objective stochastic search (MPMOSS) algorithm for generating heterogeneous excitatory and inhibitory neuronal populations

To generate physiologically validated and heterogeneous populations of cortical excitatory and inhibitory neurons, we employed the multi-parametric multi-objective stochastic search (MPMOSS) algorithm (Foster et al., 1993; Prinz et al., 2003; Marder and Taylor, 2011a; Rathour and Narayanan, 2012; Mittal and Narayanan, 2018; Mishra and Narayanan, 2019; Gorur-Shandilya et al., 2022; Sinha and Narayanan, 2022; Saini and Narayanan, 2025). This approach enabled systematic and wide exploration of the parametric space of intrinsic conductances as well as validation against subtype-specific physiological constraints, together ensuring that the resulting model population spanned a broad and biologically plausible range.

For the excitatory neuronal population, we generated 8,500 random models by independently sampling each of the twelve model parameters from predefined uniform distributions (Table 1). This random selection produced neurons with distinct combinations of intrinsic parameters. Each generated model was then validated against a set of 10 electrophysiological measurements characteristic of L5 pyramidal neurons (Table 2). A model was considered valid only if it satisfied all electrophysiological bounds together, ensuring that the final population exhibited physiologically realistic and experimentally consistent behavior across multiple functional dimensions. We applied a similar but independent MPMOSS procedure to generate the inhibitory neuronal population of models. Here, we generated 2,000 random models from a six-dimensional parametric space (Table 3) and validated them against 10 electrophysiological measurements from cortical basket cells (Table 4).

To characterize the validated excitatory and inhibitory model populations, we independently examined heterogeneities in their parametric and measurements spaces. We computed pairwise correlations among intrinsic measurements of the two valid model populations independently, allowing us to determine whether different measurements captured independent aspects of neuronal behavior or were interdependent. We analyzed pairwise correlations among the parameters underlying the two valid-model populations independently, to assess if the physiologically validated models required tightly constrained parametric interactions.

### Feedforward network architecture

We constructed feedforward networks of these excitatory and inhibitory neurons composed of well-defined sequential assemblies. The network consisted of ten assemblies, each containing 100 excitatory and 25 inhibitory neurons, consistent with cortical distribution of excitatory vs. inhibitory neurons (Beaulieu, 1993; Markram et al., 2004; Buzsaki et al., 2007). Connectivity within each assembly was probabilistic. Excitatory-to-inhibitory connections were formed with a probability of 0.06, inhibitory-to-excitatory connections with 0.373, and inhibitory-to-inhibitory connections with 0.316. As synfire chains represent a temporal sequence of feedforward propagation across the excitatory neuronal population (Diesmann et al., 1999), excitatory-to-excitatory connections within an assembly were absent. Feedforward connectivity across assemblies was fully determined: every excitatory neuron in a given assembly projected to all excitatory and inhibitory neurons in the immediate downstream assembly. This ensured that synchronous spiking in one layer could reliably influence the next, allowing for controlled propagation of activity along the chain. There were no feedback connections as the network architecture was strictly feedforward. The first assembly received 100 external inputs, which together constituted input packets which could contain a maximum of 100 spikes with a parametrized temporal spread. As with the other downstream assemblies, these inputs were fully connected to both excitatory and inhibitory neurons within the first assembly. This network design allowed us to systematically examine how intrinsic excitability, heterogeneity, and changes to ionic conductances shape the sustenance or breakdown of synchrony as it propagates across a sequence of cortical assemblies.

### Background synaptic inputs and high-conductance state

To replicate the high-conductance state that is observed *in vivo*, each neuron in the network received 5000 background synaptic inputs receiving independently random spike trains generated using a Poisson process (Destexhe et al., 2003; Mishra and Narayanan, 2015). Of these 5000 inputs, 80% were excitatory and 20% were inhibitory. The synaptic weights of these background inputs were adjusted such that a unitary excitatory postsynaptic potential (uEPSP) produced a depolarization of 0.3 mV, while a unitary inhibitory postsynaptic potential (uIPSP) produced a hyperpolarization of 1 mV (Stuart, 1999; Chance et al., 2002; Holmgren et al., 2003; Kubota et al., 2016). Excitatory background inputs arrived at an average rate of 2 Hz per synapse, and inhibitory inputs were delivered at 2.2 Hz per synapse. These rates were chosen to match the number of synaptic inputs and their unitary potentials, so that the average membrane potential of the neuron remained largely unperturbed after the onset of background synaptic inputs, thereby maintaining excitation-inhibition balance. We assessed the impact of a high-conductance state by measuring the neuron’s input resistance (*R*_*in*_measured using the slope of the steady-state *V* − *I* plot) before and after background inputs were introduced. Specifically, we injected current steps ranging from −50 pA to +50 pA in steps of 25 pA. This was repeated 10 times for each current value, each time with a different random seed governing the background synaptic inputs. For each current injection, we averaged the voltage responses across the 10 trials and then computed the input resistance from the average steady-state voltage deflection.

### Input packets

All neurons in the first assembly of the synfire chain received 100 excitatory synaptic inputs. The nature of spike trains that were fed to these 100 inputs were parametrized by *a*_*in*_, representing the number of spikes in the input packet, and *σ*_*in*_, denoting the temporal spread of spikes in the packet (Diesmann et al., 1999). For every analyzed network, propagation across all 10 assemblies was tested for a total of 340 input combinations, involving 20 values of *a*_*in*_ (from 5 to 100 in steps of 5) and 17 values of *σ*_*in*_ (from 0 to 4 ms in increments of 0.25 ms). For any (*a*_*in*_, *σ*_*in*_) combination, *a*_*in*_ number of input spike timings were sampled from a Gaussian distribution with a fixed mean at 700 ms and standard deviation *σ*_*in*_. Propagation of each input packet was assessed over a 1000-ms simulation period, with the input packet introduced at 700 ms to allow time for stabilization of all conductance-based models. The simulation period spanning from ∼670 ms to 800 ms was considered for all analyses. For each combination of these input parameters, we ran 10 independent trials, together yielding 3,400 propagation simulations across all 10 assemblies for each network configuration. Each trial differed in the pattern of background synaptic inputs that neurons received, thereby incorporating variability in the network’s ongoing activity state. The input packets were resampled from the corresponding Gaussian distribution for each independent trial.

### Varying HCN-channel densities and computation of spike triggered average

To study how HCN channels influence the propagation of synchrony, we scaled HCN-channel conductance values across the population of neurons in the network. Specifically, we studied the impact of increasing or decreasing the HCN channel density by a factor of 8 relative to a baseline condition, resulting in 3 HCN channel densities: low HCN (1/8×), baseline HCN (1×), and high HCN (8×). To quantify how these changes in HCN conductance altered intrinsic properties, we computed neuronal intrinsic properties at the three densities of HCN channels. In addition, to assess spike initiation dynamics as a function of HCN-channel density, we computed the spike-triggered average (STA) of neurons under each of these three conditions (Aguera y Arcas and Fairhall, 2003; Hong et al., 2007; Das and Narayanan, 2014, 2015, 2017; Mittal and Narayanan, 2018).

To compute STA, we injected zero-mean Gaussian white noise current into a model neuron. The standard deviation of the injected noise was adjusted so that the neuron fired at an average inter-spike interval (ISI) of approximately one second. This level of noise ensured that spikes occurred with sufficient regularity, while preserving variability in the input current to capture meaningful features in the STA (Aguera y Arcas and Fairhall, 2003; Hong et al., 2007; Das and Narayanan, 2014, 2015, 2017; Mittal and Narayanan, 2018). We recorded a total of 2000 spike events that were separated by at least one second from each other, ensuring statistical independence between spikes that were sampled. For each of these spikes, we extracted 200 ms of the injected noise current trace that preceded the spike and saved that segment. The spike-triggered average was then computed by averaging the input current segments across all spikes. This provided a time-resolved profile of the average input preceding a spike and served as a proxy for the effective input filter of the neuron under different HCN densities. This procedure was repeated for all three HCN conditions, baseline, 8×, and 1/8×, to compare how the temporal structure of inputs leading to spike generation changed as a function of intrinsic membrane conductance.

### Heterogeneous network of feedforward assemblies

To explore how within-type intrinsic heterogeneity affects propagation of synchrony in a synfire chain network, we simulated multiple versions of the network with varying degrees of intrinsic heterogeneity spanning both excitatory and inhibitory neuronal populations. Each degree of heterogeneity was defined by the number of unique neuronal instances that were used to build any given assembly. For all of these, excitatory and inhibitory neurons were selected at random from the respective heterogeneous populations validated using MPMOSS.

Degree 1 represented the lowest degree of heterogeneity, whereby each assembly was internally homogeneous, built of repeating homogeneous units (one excitatory and one inhibitory neuron). To be specific, a single excitatory neuronal model was repeated to make 100 excitatory neurons in the assembly and the 25 inhibitory neurons were repeated from a single inhibitory neuronal model. At Degree 2, each assembly was constructed from two randomly selected excitatory neurons and two random inhibitory neurons, each distributed randomly to form 100 excitatory and 25 inhibitory neurons in the assembly. Each of the 10 assemblies, however, were built with distinct different excitatory and inhibitory neuronal pairs. Intermediately higher degrees of heterogeneity followed the same procedure, with four, eight, sixteen, thirty-two, or sixty-four excitatory and inhibitory neurons per assembly at Degrees 4, 8, 16, 32, and 64, respectively. At the highest degree of heterogeneity, Degree 100, every single neuron in the assembly was unique and picked randomly from the respective excitatory or inhibitory neuronal population. Across all degrees of heterogeneity, each assembly was constructed with distinct sets of excitatory and inhibitory neurons while respecting the degree of heterogeneity across all assemblies. This experimental design ensured that we assessed synchrony propagation across a graded spectrum of heterogeneity degrees, ranging from a totally homogeneous assembly at Degree 1 to fully heterogeneous assemblies built of unique neurons at Degree 100. Note that the numbering in the nomenclature associated with the degrees is with reference to the number of distinct excitatory neurons. While the number of unique inhibitory neurons progressively increased from 1 to 16 through heterogeneity Degrees 1–16 (as with the unique excitatory neurons), the inhibitory population was completely heterogeneous from Degree 32 onward as there were only 25 inhibitory neurons in each assembly.

As neuronal models were picked randomly to instantiate networks, we generated 12 independent network instantiations for each degree of heterogeneity. In each case, different neurons were randomly selected from the full pool to populate each assembly, while adhering to number picked for each degree of heterogeneity. The use of a population of networks to study the impact of heterogeneities (i) ensured that the observed effects were not dependent on the specific identity of the randomly picked neurons; (ii) allowed us to assess network-to-network variability in propagation efficacy; and (iii) allowed for the possibility of arriving at generalizable consequences of heterogeneity that are observed across networks (Prinz et al., 2004; Schaeffer et al., 2024; Saini and Narayanan, 2025; Santhosh and Narayanan, 2025, 2026). As before, for each network, we studied propagation using input packets composed of 340 different combinations of *a*_*in*_ and *σ*_*in*_, with 10 trials for each combination. Together, a total of 12 independent networks, each initialized with 8 degrees of heterogeneities and 3 levels of HCN were studied with 10 trials of 340 different input combinations, yielding a total of 979,200 (12 × 8 × 3 × 10 × 340) network simulations studying propagation across all assemblies.

An additional set of simulations were performed with fast HCN channels, where we reduced the activation time constant 𝜏_*HCN*_ by a factor of twenty, which effectively removed the slow feedback dynamics that normally accompany HCN activation and yielded a fast-HCN channel (Narayanan and Johnston, 2007, 2008; Das and Narayanan, 2014; Sinha and Narayanan, 2015; Mishra and Narayanan, 2025). We replaced the endogenous slow HCN channels in all neurons in the network with their fast counterparts, matched their excitability profiles by adjusting the fast HCN conductance value to match maximal impedance amplitudes (Das and Narayanan, 2014; Sinha and Narayanan, 2015), and performed an additional set of network simulations. Specifically, a further set of 12 × 3 × 10 × 340 = 122,400 simulations were performed with fast HCN channels for 12 independent homogeneous (Degree 1) networks, each initialized at 3 levels of fast HCN conductances studied with 10 trials of 340 different input combinations. Thus, the total number of feedforward propagations through all 10 assemblies used in this study were 1,101,600.

### Quantification of synchrony propagation efficacy

To quantitatively compare synchrony propagation in different networks with several graded degrees of heterogeneity and different HCN-channel densities and kinetics, we defined propagation efficacy as the average probability that an input volley successfully propagated through all assemblies of the feedforward network. To compute this, we first calculated the probability of successful propagation across 10 trials, for each of 340 input combinations. For any given input combination, propagation was defined to be successful if the packet size in the last assembly (Assembly 10) was greater than 40. For a given input combination *i*, let *N*_*prop*_(*i*) represent the number of trials (of the total 10 trials) where successful propagation was observed when input combination *i* was presented to the network. Then, the probability of propagation 𝑝_*prop*_(*i*) of any input combination *i* for that network was defined as the ratio between *N*_*prop*_(*i*) and the total number of trials. Synchrony propagation efficacy (𝐸_*prop*_) was then defined as the average probability of propagation across all 340 input combinations:

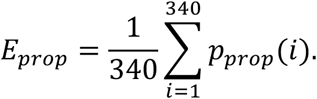

As every stimulus combination was fully represented by the associated (*a*_*in*_, *σ*_*in*_) values, the 340 values of 𝑝_*prop*_(*i*) were visualized as a matrix of 20 rows of *a*_*in*_ (from 5 to 100 in steps of 5) and 17 columns of *σ*_*in*_(from 0 to 4 ms in increments of 0.25 ms). By definition, 0 ≤ 𝐸_*prop*_ ≤ 1, with 𝐸_*prop*_ = 1 representing a scenario where the network was able to successfully propagate synchrony for all 340 input combinations, while 𝐸_*prop*_ = 0 indicated the inability of the network to propagate any input combination. 𝐸_*prop*_was used as a measure of propagation efficacy of different networks with several graded degrees of heterogeneity and different HCN-channel densities and kinetics.

We defined 𝑊_II_ as the width of the stochastic separatrix that separated the completely successful (𝑝_*prop*_(*i*) = 1) and completely failed (𝑝_*prop*_(*i*) = 0) input combinations. 𝑊_II_was calculated as the number of input combinations where 0 < 𝑝_*prop*_(*i*) < 1 across all 340 input combinations:

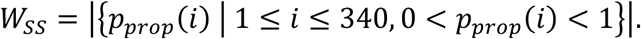

Packet size and spread at the fixed points of convergence for successfully propagating inputs for a given network were computed as the average output packet size (*a*_𝑜𝑢𝑡_) and spread (*σ*_𝑜𝑢𝑡_) of the final (10^th^) assembly, averaged across all successfully propagating inputs (across trials) for the network under consideration.

### Computational details

All simulations were performed using the NEURON simulation environment, with a fixed *dt* of 0.025 ms and at a temperature of 34° C. All data analysis and visualization were carried out using MATLAB. All statistical analyses were performed using R.

## RESULTS

Information propagates across the brain in noisy environments. This information can be encoded either in the rate of action potentials or in their precise timing. For timing-based coding, it is critical that neuronal circuits have mechanisms to generate precisely timed spikes and to reliably transmit them to downstream targets. Previous studies have shown that neurons can generate precisely timed action potentials, especially for stimulus-driven spiking (O’Keefe and Recce, 1993; Bair and Koch, 1996; Mehta et al., 2002; Ariav et al., 2003; Jensen, 2005; Maimon and Assad, 2009; Buzsaki, 2010; Churchland et al., 2010; Ratte et al., 2013; Levenstein et al., 2026). Several computational and experimental studies have examined how input statistics and network architecture could influence the propagation of synchronous activity (Ermentrout, 1996; Diesmann et al., 1999; Reyes, 2003; Abouzeid and Ermentrout, 2009; Kremkow et al., 2010; Kumar et al., 2010; Hong et al., 2012; Jahnke et al., 2013; Ratte et al., 2013; Moldakarimov et al., 2015). However, the impact of neuronal biophysical properties and heterogeneities therein on synchrony propagation across feedforward assemblies remains poorly understood. In this expansive study, we assess synchrony propagation across a sequence of conductance-based heterogeneous assemblies constructed with excitatory and inhibitory neurons (Fig. 1). We systematically evaluate synchrony propagation in a large population of feedforward networks endowed with different degrees of heterogeneity and distinct ion-channel combinations.

### Heterogeneous populations of cortical excitatory and inhibitory neurons

As with all neuronal subtypes, cortical excitatory and inhibitory neurons manifest within-type heterogeneities in molecular composition and physiology (Shamir and Sompolinsky, 2006; Padmanabhan and Urban, 2010; Hay et al., 2011; Tripathy et al., 2013; Cembrowski and Spruston, 2019; Tzilivaki et al., 2019; Moradi Chameh et al., 2021; Rich et al., 2022; Planert et al., 2025; Dahmen et al., 2026). In assessing the impact of within-type heterogeneities on circuit function, the first step was to generate heterogeneous populations of cortical excitatory and inhibitory neurons in a manner that they reflect subtype-specific characteristics and heterogeneity therein. As our specific question focused on biophysical characteristics of neurons, we built conductance-based models of cortical excitatory and inhibitory neurons (Fig. 1*A*). The set of ion channels expressed in these models and their gating kinetics were subtype-specific and derived from previous computational models (Hay et al., 2011; Tzilivaki et al., 2019).

To generate heterogeneous population of both neuronal subtypes, we employed two independent instances of the well-established multi-parametric multi-objective stochastic search (MPMOSS) algorithm (Foster et al., 1993; Prinz et al., 2003; Marder and Taylor, 2011a; Rathour and Narayanan, 2012; Mittal and Narayanan, 2018; Mishra and Narayanan, 2019; Gorur-Shandilya et al., 2022; Sinha and Narayanan, 2022; Saini and Narayanan, 2025). Specifically, for each neuronal subtype, we first constructed a hand-tuned base model by iteratively adjusting subtype-specific parameters until the model reproduced the characteristic electrophysiological signatures of that subtype. These base models served as substrates for defining parameter ranges in a subsequent large-scale stochastic search, involving a broad and physiologically plausible parametric space. We generated thousands of candidate models for each subtype, by randomly sampling their parametric space, and computed electrophysiological measurements for each model. We compared these model measurements against the corresponding subtype-specific signatures, and models were declared valid only if they satisfied all specified electrophysiological criteria.

For the excitatory neuronal population, we generated 8,500 random single-neuron models by sampling from a 12-dimensional parametric space, with each parameter drawn independently from its corresponding uniform distribution (Table 1). All models were validated against a set of 10 electrophysiological measurements characteristic of layer 5 pyramidal neurons (Table 2) and 224 models (∼2.6%) were found to be valid. We found that the measurements (Fig. 2*A*) and parameters (Fig. 2*B*) of these valid excitatory neuronal models spanned their respective permissible ranges (Tables 1–2), indicating that the resulting population was heterogeneous in the measurement as well as in the parametric space. For the inhibitory population, we sampled a six-dimensional parametric space (Table 3) for generating 2,000 random single-neuron models and validated them against 10 electrophysiological measurements derived from cortical basket cells (Table 4). We found 124 (∼6.2%) of these models to be valid, with measurements (Fig. 3*A*) and parameters (Fig. 3*B*) manifesting heterogeneous distributions spanning their respective valid ranges (Tables 3–4).

**Figure 2:**
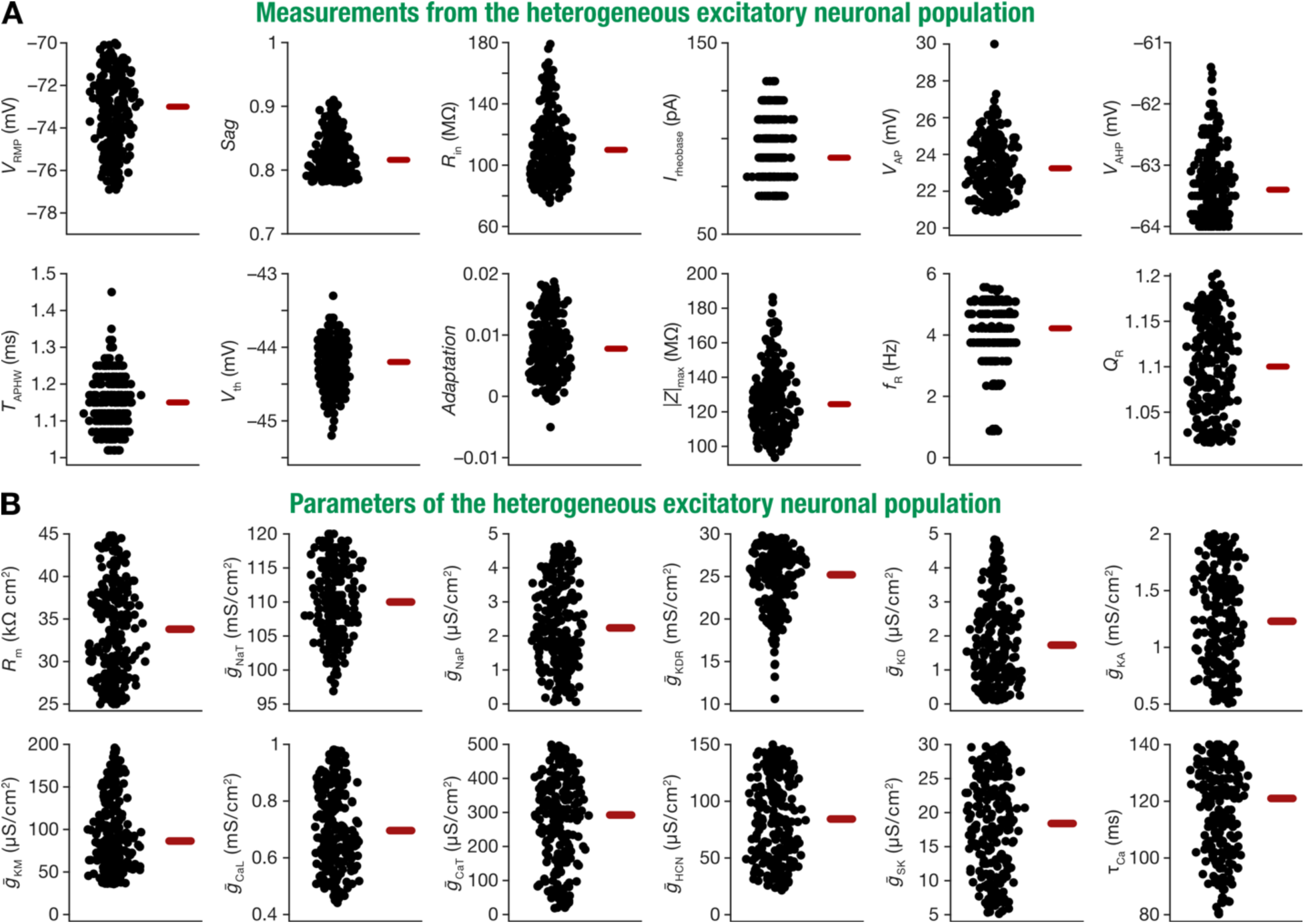
Within-type heterogeneities in measurements and parameters of the excitatory neuronal population. (A) Beeswarm plots of intrinsic electrophysiological measurements of all 224 valid excitatory neurons generated using MPMOSS. All measurements spanned the entire range defined for their validation (Table 2). (B) Beeswarm plots of model parameters for all 224 valid excitatory neurons generated using MPMOSS, plotted to span the lower-to-upper bounds of each parameter (Table 1). All thick red lines represent the respective median values.

**Figure 3:**
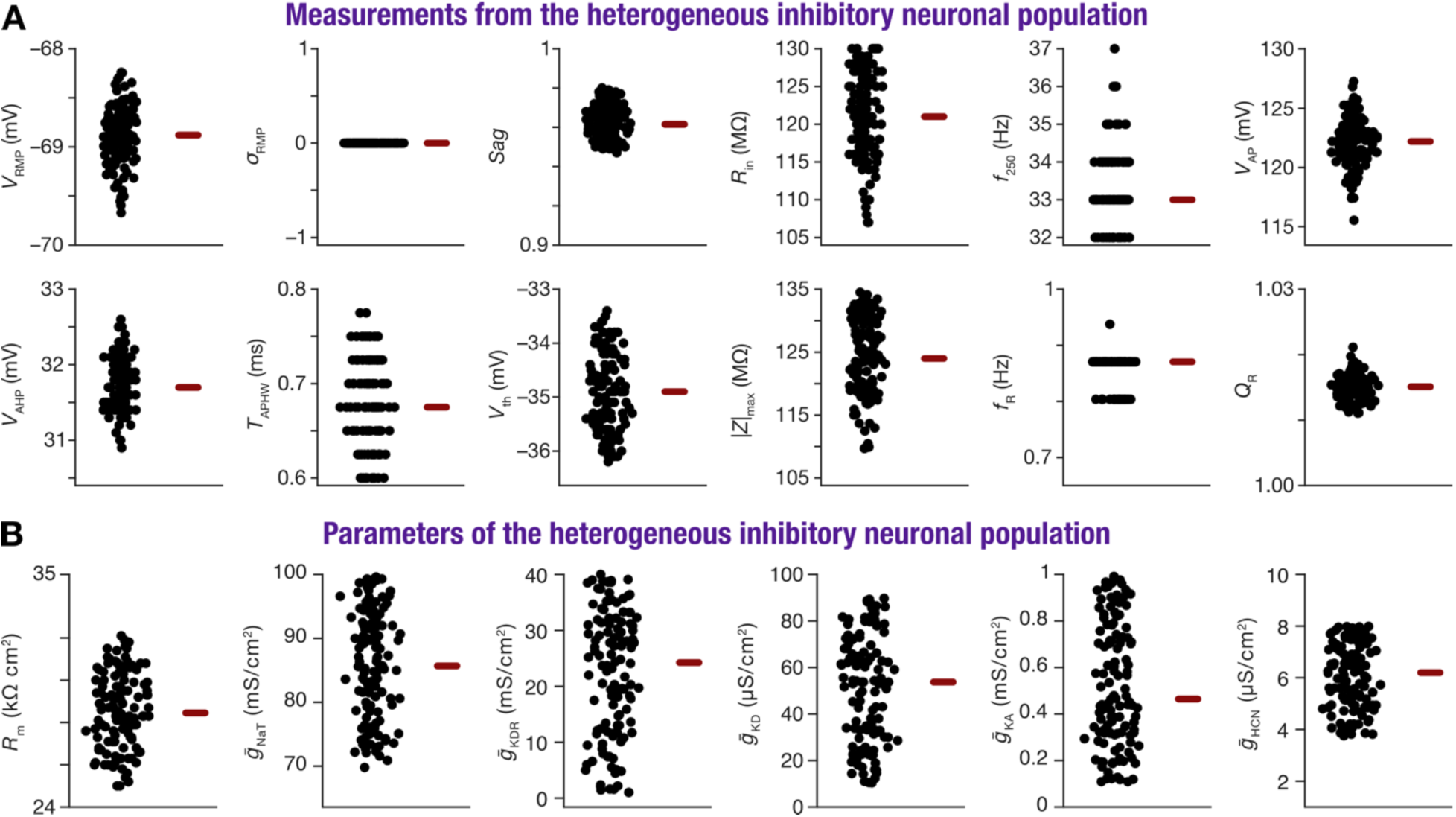
Within-type heterogeneities in measurements and parameters of the inhibitory neuronal population. (A) Beeswarm plots of intrinsic electrophysiological measurements of all 124 valid inhibitory neurons generated using MPMOSS. All measurements spanned the entire range defined for their validation (Table 4). (B) Beeswarm plots of model parameters for all 124 valid inhibitory neurons generated using MPMOSS, plotted to span the lower-to-upper bounds of each parameter (Table 3). All thick red lines represent the respective median values.

We computed pairwise correlations across the different electrophysiological measurements of valid models to assess if they were indeed characterizing different aspects of neuronal physiology. For the valid excitatory neuronal population, the correlations were weak for most measurement pairs, except for measurements where strong relationships were expected (Fig. 4*A*).

**Figure 4:**
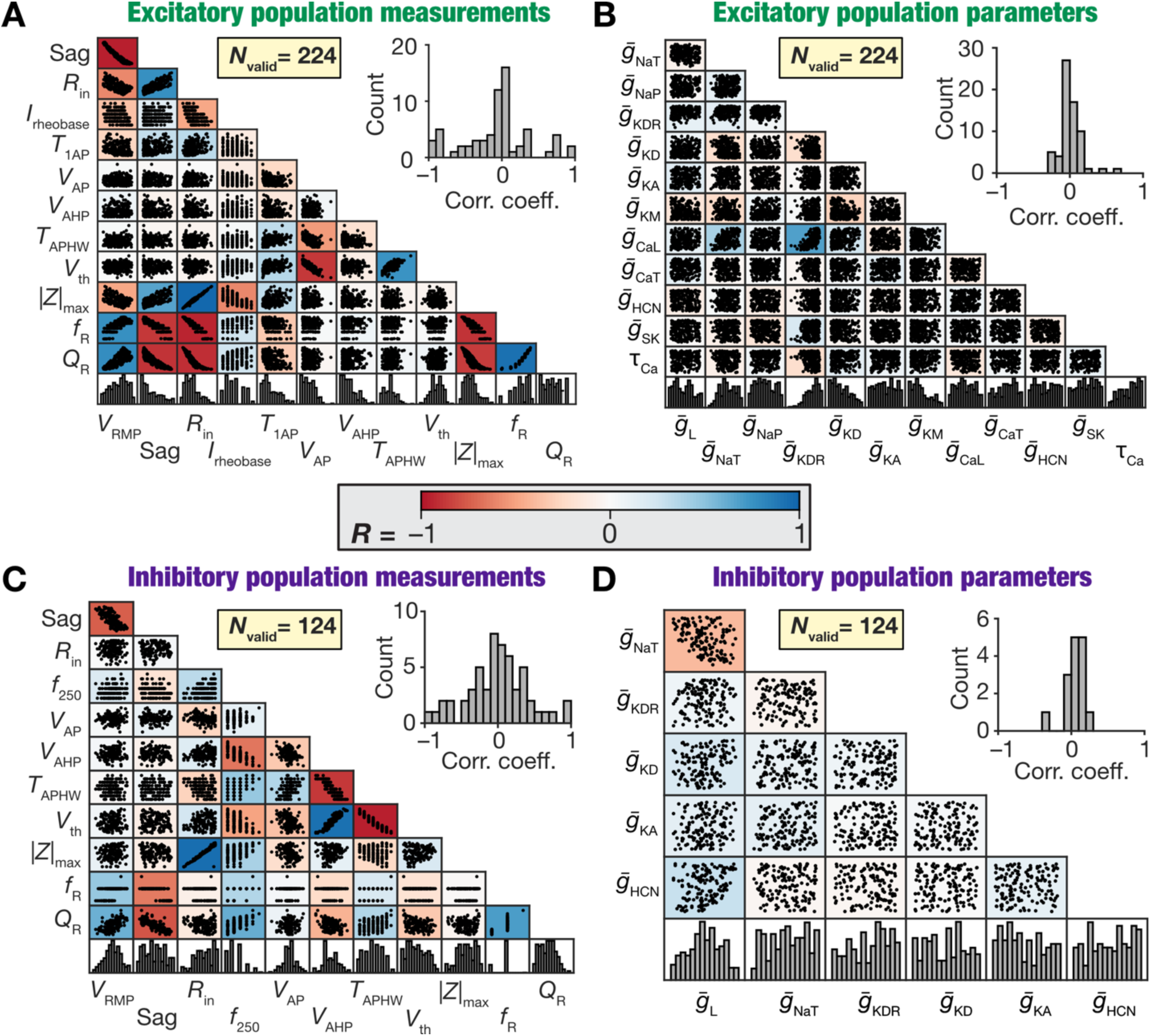
Pairwise relationships in measurements and parameters of the heterogeneous conductance-based excitatory and inhibitory neuronal model populations. (A–B) Scatter plot matrix of measurements (A) and parameters (B) from the valid population of excitatory neuronal models obtained using MPMOSS. (C–D) Scatter plot matrix of measurements (C) and parameters (D) from the valid population of inhibitory neuronal models obtained using MPMOSS. The bottom-most rows in each panel depict the histogram of the specific measurement or parameter in that column, illustrating widespread heterogeneity across models. The insets within each panel show the histogram of the Pearson’s correlation coefficient values, which are overlaid on the respective scatter plot matrices, for the unique correlation coefficient values from the pairwise comparisons shown in the scatter plot matrix.

For instance, we found strong relationships among resting membrane potential, input resistance, sag ratio, maximum impedance, resonance frequency, and resonance strength, which was expected given these measurements covary due to their shared dependence on subthreshold conductances such as HCN channels (Robinson and Siegelbaum, 2003; Narayanan and Johnston, 2007, 2008; Combe and Gasparini, 2021; Mishra and Narayanan, 2025). Importantly, Pearson correlation coefficients computed across parameters of valid excitatory neurons were weak, showing that strong pairwise dependencies among parameters were not required to yield signature electrophysiological properties (Fig. 4*B*). We found similarly weak pairwise relationships between inhibitory neuronal measurements (Fig. 4*C*), except for measurements where strong relationships were expected, and weak pairwise relationships between the associated parameters (Fig. 4*D*).

Together, the MPMOSS procedures yielded heterogeneous valid model populations that captured the experimentally observed variability in the intrinsic properties of cortical excitatory and inhibitory neurons. These physiologically validated heterogeneous populations were used to build heterogeneous feedforward networks, ensuring that all network-level analyses were grounded in biologically realistic and non-repeating neuronal compositions. Importantly, these analyses demonstrated that several non-random (as most random models were invalid) and non-unique (as the underlying parametric distributions across are broad) combinations of functionally specialized molecular subsystems (ion channels, calcium buffers) yielded signature cellular function. These observations are consistent with reduced models of other neuronal subtypes where such complexity and degeneracy (the ability of disparate structural components to yield similar function) have been demonstrated (Prinz et al., 2004; Taylor et al., 2009; Marder and Taylor, 2011b; Rathour and Narayanan, 2012; Srikanth and Narayanan, 2015; Mittal and Narayanan, 2018; Mishra and Narayanan, 2019; Mittal and Narayanan, 2022; Yang et al., 2022; Schneider et al., 2023).

### High-conductance state regulated neuronal intrinsic properties

The dependencies of neuronal intrinsic properties on high-conductance state (Chance et al., 2002; Destexhe et al., 2003; Mishra and Narayanan, 2015) are well-established. High-conductance state, generated by background synaptic activity that activate excitatory and inhibitory synaptic receptors, is an important distinguishing feature of *in vivo* neural circuits (Destexhe et al., 2003). We introduced high-conductance state into our neuronal models by having each of them receive 5,000 randomly distributed background synaptic inputs, composed of 80 percent excitatory and 20 percent inhibitory synapses (Mishra and Narayanan, 2015). The presynaptic activation rates of these background inputs were tuned to maintain a balanced state between excitation and inhibition, with the unitary excitatory and inhibitory postsynaptic potential set to 0.3 mV and 1 mV, respectively (Supplementary Fig. S1*A*). We verified this balance by monitoring average resting membrane potential before and after background synaptic activity was introduced. The membrane potentials remained stable, confirming that network activity was balanced and did not drift toward either excitation or inhibition (Supplementary Fig. S1*B*). As all synapses in the network were modeled using conductance-based formulations, elevated background synaptic activity automatically translated to an increase in total membrane conductance, together resulting in a high-conductance state. Therefore, the introduction of background synaptic activity expectedly (Chance et al., 2002; Destexhe et al., 2003; Mishra and Narayanan, 2015) reduced the input resistance of excitatory and inhibitory neurons (Supplementary Fig. S1*C–E*). The significant reduction in input resistance confirmed that the neuron had transitioned into a high-conductance regime, consistent with experimental observations in active cortical circuits (Destexhe et al., 2003).

In addition to these background synaptic inputs, each neuron in every assembly also received two additional spike trains, one excitatory and one inhibitory, with stronger synaptic weights. These spike trains were randomly timed, with excitatory and inhibitory inputs arriving at a frequency of 2 Hz and 2.2 Hz, respectively. The large excitatory inputs generated spontaneous firing in both excitatory and inhibitory neurons (Supplementary Fig. S1*F*) and the large inhibitory inputs acted to counterbalance the impact of their excitatory counterparts on the average membrane potential. Together, this input regime ensured that all neurons in the network operated within a balanced high-conductance regime and generated random spontaneous action potentials.

### Construction of feedforward network of assemblies

We constructed feedforward network models built of 10 assemblies using the valid-model populations of excitatory (Fig. 2) and inhibitory (Fig. 3) neurons. In building up to assessing the impact of within-type heterogeneities on synchrony propagation, we first examined homogeneous synfire chain networks where each assembly was built with a single repeating homogeneous unit of an excitatory and an inhibitory neuron. Specifically, a single excitatory neuronal model was repeated to make 100 excitatory neurons in the assembly and the 25 inhibitory neurons were repeated from a single inhibitory neuronal model. Each of the 10 assemblies, however, were built with distinct neuronal pairs. The first assembly received an input packet composed of 100 synaptic inputs that projected to all excitatory and inhibitory neurons in the assembly. The input packet was parametrized by *a*_*in*_, representing the number of spikes in the input packet, and *σ*_*in*_, denoting the temporal spread of spikes in the packet (Diesmann et al., 1999). For every analyzed network, propagation across all 10 assemblies was tested for a total of 340 input combinations, involving 20 values of *a*_*in*_ (from 5 to 100 in steps of 5) and 17 values of *σ*_*in*_ (from 0 to 4 ms in increments of 0.25 ms). For any (*a*_*in*_, *σ*_*in*_) combination, *a*_*in*_ number of input spike timings were sampled from a Gaussian distribution with a fixed mean and standard deviation *σ*_*in*_ .

Within each assembly, connectivity among neurons was probabilistic. The connection probabilities were 0.06 for E-to-I, 0.373 for I-to-E, 0.316 for I-to-I, and 0 for E-to-E connections. Synaptic strengths were adjusted such that individual excitatory inputs generated excitatory postsynaptic potentials (EPSPs) of approximately 0.3 mV, while inhibitory inputs produced inhibitory postsynaptic potentials (IPSPs) of approximately 1 mV. Thus, each neuron in every assembly received feedforward inputs from the previous assembly (or the input packet for the first assembly), within assembly inputs from the other neurons in the assembly, and background synaptic inputs that governed high-conductance states and spontaneous firing. This configuration provided the foundation for studying how input activity patterns introduced into the first assembly propagated through successive assemblies and how intrinsic neuronal properties shaped the fidelity of this propagation.

### Stochastic separatrix separated propagating from non-propagating input packets

Using this network configuration, we first systematically examined how synchronous activity propagates across a single homogeneous assembly of conductance-based model neurons endowed with high-conductance states. To do this, we delivered input packets with different (*a*_*in*_, *σ*_*in*_) combinations and computed (*a*_𝑜𝑢𝑡_, *σ*_𝑜𝑢𝑡_), the output packet size and spread after one assembly of propagation (Diesmann et al., 1999). Consistent with earlier observations using simple spiking models without high-conductance states (Diesmann et al., 1999), we found that output packet size increased with increasing input packet size and output packet spread was higher with larger input packet spread (Fig. 5*A*). There were cross-interactions as well, with larger input packet spread translating to lesser *a*_𝑜𝑢𝑡_ and larger input packet size resulting in lesser *σ*_𝑜𝑢𝑡_. For each pair of (*a*_*in*_, *σ*_*in*_), we assessed propagation performance across 10 independent trials, with trials differing in terms of the seed values that defined background synaptic inputs. Owing to the stochasticity associated with the background synaptic inputs, we observed pronounced trial-to-trial variability in (*a*_𝑜𝑢𝑡_, *σ*_𝑜𝑢𝑡_) across each pair of (*a*_*in*_, *σ*_*in*_) (Fig. 5*A*). Although large number of synchronous inputs were ideal for the purpose, even a small number of tightly synchronized inputs or a large number of weakly synchronous inputs were sufficient to recruit the downstream population towards synchrony propagation (Diesmann et al., 1999). Together, these observations confirmed prior observations (Diesmann et al., 1999) that small input packet sizes can yield synchronous output if their spread is low, and that synchronous outputs could still be achieved with high input packet spread if the packet size is larger.

**Figure 5:**
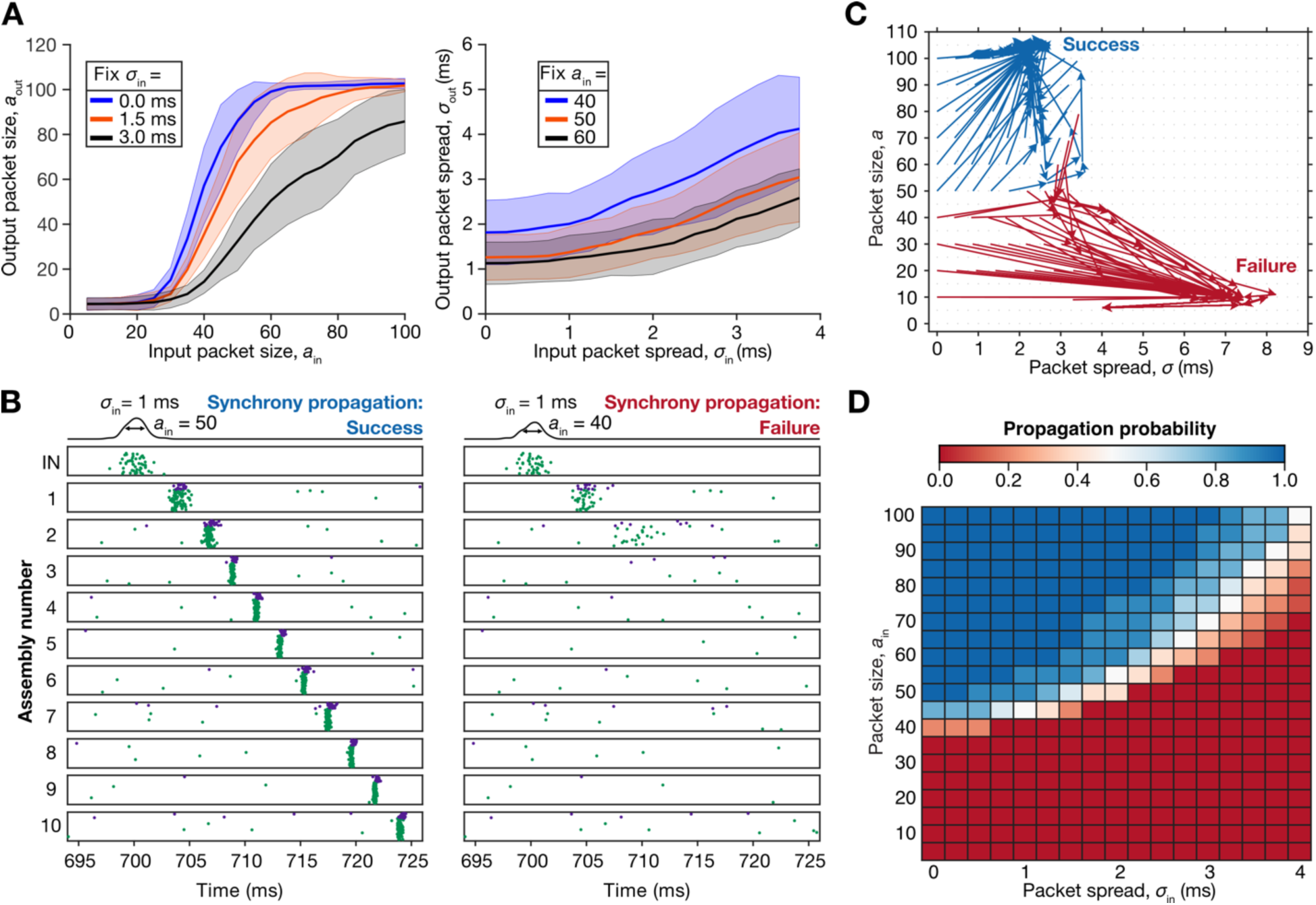
Stochastic separatrix defining success in synchrony propagation across homogeneous excitatory-inhibitory synfire chains. (A) *Left*, Output packet size (*a*_𝑜𝑢𝑡_) increases as input packet spread (*σ*_*in*_) decreases for a fixed input packet size (*a*_*in*_). *Right*, Output packet spread (*σ*_𝑜𝑢𝑡_) decreases with increasing input size (*a*_*in*_) for a fixed input packet spread (*σ*_*in*_). These plots refer to propagation across a single assembly. The shaded regions represent the standard deviation, computed by running 10 trials for each (*a*_*in*_, *σ*_*in*_) pair. (B) *Left*, Raster plots showing successful propagation of an input spike packet with *a*_*in*_ = 50 and *σ*_*in*_ = 1 ms. *Right*, Raster plots showing failure of propagation of an input spike packet with *a*_*in*_ = 40 and *σ*_*in*_ = 1 ms. Both sets of raster plots depict the outcomes of a single trial of propagation. (C) State space of volley propagation in a trial, for a representative subset of input combinations of input packet size and spread. (D) Propagation probability matrix computed across 10 trials for each input packet size (*a*_*in*_) and spread (*σ*_*in*_) highlights the emergence of a stochastic separatrix. For instance, packets with *a*_*in*_ = 10 and *σ*_*in*_ = 4 ms never propagated successfully across all 10 trials, packets with *a*_*in*_ = 90 and *σ*_*in*_ = 1 ms always showed successful synchrony propagation across all 10 trials, and packets with *a*_*in*_ = 55 and *σ*_*in*_ = 2 ms showed probabilistic propagation, with successful propagation for half the trials and failure for the other half. The propagation efficacy 𝐸_*prop*_ for this network was computed to be 0.37 and the width of the stochastic separatrix 𝑊_𝑆𝑆_ was 72.

To examine how input packets propagated across the feedforward network, we computed *a*_𝑜𝑢𝑡_and *σ*_𝑜𝑢𝑡_after each assembly as specific input packets propagated across all 10 assemblies. Two different (*a*_*in*_, *σ*_*in*_) combinations, with one resulting in successful synchrony propagation and another failing to propagate are shown in Figure 5*B*. We noted a transient period of suppression of spontaneous spikes after strong synchrony in any assembly, owing to strong inhibitory projections suppressing firing (Fig. 5*B*). Representing (*σ*, *a*) combinations on a phase plane (Diesmann et al., 1999), we plotted (*σ*, *a*) as a given (*a*_*in*_, *σ*_*in*_) propagated across every assembly for a single trial (Fig. 5*C*). Specifically, for each (*a*_*in*_, *σ*_*in*_), 10 connected lines with arrows depict (*σ*, *a*) through the propagation across all assemblies. The first line starts at (*σ*_*in*_, *a*_*in*_) and ends with an arrow at (*σ*_𝑜𝑢𝑡_, *a*_𝑜𝑢𝑡_) of the first assembly. The second line starts at the (*σ*_𝑜𝑢𝑡_, *a*_𝑜𝑢𝑡_) of the first assembly and ends with an arrow at (*σ*_𝑜𝑢𝑡_, *a*_𝑜𝑢𝑡_) of the second assembly, with this continuing through the 10 assemblies (Fig. 5*C*). The phase-plane representation visually confirmed earlier observations that successful propagation occurred with a conjunctive threshold on the minimum *a*_*in*_ and maximum *σ*_*in*_ that will allow propagation (Fig. 5*C*). In addition, the phase plane depicted the existence of a boundary separating propagating from non-propagating combinations, which previous studies have referred to as a separatrix (Diesmann et al., 1999).

These representations were limited to a single trial of propagation for each of the 340 combinations of (*a*_*in*_, *σ*_*in*_). To assess trial-to-trial variability, as before (Fig. 5*A*), we assessed propagation spanning all assemblies, across ten trials for each combination of (*a*_*in*_, *σ*_*in*_). Strikingly, we found that the separatrix does not form the exact same strict boundary between propagating and non-propagating regimes across different trials but showed trial-to-trial variability. This was consequent to scenarios where the same input packet propagated successfully in some trials but failed to propagate in others. To systematically quantify this trial-to-trial variability in propagation, we computed propagation probability (𝑝_*prop*_) across the 10 different trials for each input combination as the proportion of trials where successful propagation occurred. We plotted propagation probability as functions of (*a*_*in*_, *σ*_*in*_) and found a narrow zone of uncertainty where 0 < 𝑝_*prop*_ < 1, representing regions where propagation was trial-dependent (Fig. 5*D*). We refer to this zone of uncertainty that separated regions where synchrony propagation always occurred (𝑝_*prop*_ = 1) from regions where synchrony propagation never occurred (𝑝_*prop*_ = 0) as the stochastic separatrix (Fig. 5*D*). As network and afferent input characteristics remained identical across trials, this variability was due to trial-to-trial fluctuations in background synaptic activity. Thus, for input combinations residing within the stochastic separatrix, even small differences in the timing or magnitude of background synaptic events can strongly alter synchrony propagation. To quantitatively assess the characteristics of this stochastic separatrix in networks of neurons with different degrees of heterogeneities and different intrinsic properties, we computed two measurements. The first measurement was propagation efficacy (𝐸_*prop*_) defined as the average probability of propagation across all 340 input combinations. By definition, 𝐸_*prop*_ would be high for networks where there were more input combinations with 𝑝_*prop*_ = 1, and low if more input combinations had low propagation probability. While 𝐸_*prop*_ quantified the global position of the stochastic separatrix, more granularly, we noticed propagation efficacy decreased as a function of *σ*_*in*_, with more *a*_*in*_ combinations showing 𝑝_*prop*_ = 1 for lower values of *σ*_*in*_. Consequently, the values of *a*_*in*_ *within* the stochastic separatrix monotonically increased as a function of *σ*_*in*_ (Fig. 5*D*), further emphasizing the codependence of successful synchrony propagation on *a*_*in*_ and *σ*_*in*_. The second measurement, the width of the stochastic separatrix 𝑊_𝑆𝑆_, quantified the extent of the zone of uncertainty and was quantitatively defined as the number of input combinations where 0 < 𝑝_*prop*_ < 1. While 𝑊_𝑆𝑆_ quantified the overall width of the stochastic separatrix, more granularly, we noticed the width of the stochastic separatrix to increase as a function of *σ*_*in*_demonstrating an increase in trial-to-trial variability with increasing input packet spread (Fig. 5*D*). Together, our results demonstrate pronounced trial-to-trial variability in the propagation success of certain input combinations, which manifested as a stochastic separatrix that formed a zone of uncertainty between the propagating and non-propagating regions of the different input combinations. From a more granular standpoint, propagation efficacy decreased and the width of the stochastic separatrix increased with input packet spread, indicating a strong dependence of trial-to-trial variability on synchrony in the input packet.

### HCN channels enhance synchrony propagation in homogeneous feedforward networks

Intrinsic properties of neurons can be modified through developmental processes, neuromodulation, activity-dependent plasticity, or pathological conditions. These provide dynamic routes to alter how neurons process inputs and could consequently regulate how synchrony is transmitted through neural circuits (Hong et al., 2012; Ratte et al., 2013). Among the several ion channels that are known to modulate neuronal intrinsic properties, the hyperpolarization-activated cyclic nucleotide-gated (HCN) channel is an attractive candidate for several reasons. The dependencies of neuronal intrinsic properties on HCN channels (Gasparini and DiFrancesco, 1997; Magee, 1998; Williams and Stuart, 2000; Robinson and Siegelbaum, 2003; Narayanan and Johnston, 2007, 2008; Shah, 2014; Combe and Gasparini, 2021; Mishra and Narayanan, 2025) are well-established. As HCN channel density can be altered by neuromodulation or activity-dependent plasticity (Pape, 1996; Fan et al., 2005; Rosenkranz and Johnston, 2006; Brager and Johnston, 2007; Narayanan and Johnston, 2007; Rosenkranz and Johnston, 2007; Narayanan and Johnston, 2008; Dembrow et al., 2010; Dembrow and Johnston, 2014; Mishra and Narayanan, 2025), we assessed the impact of scaling HCN-channel density on neuronal intrinsic properties. We confirmed that scaling HCN-channel density altered sub- and supra-threshold properties along expected lines in our model, specifically depolarizing resting membrane potential, reducing input resistance, and increasing resonance frequency (Supplementary Fig. S2*A–E*). As expected from previous literature (Mishra and Narayanan, 2015), the strong depolarization introduced by increasing HCN-channel density was strongly suppressed in the presence of high-conductance state, although the reduction in input resistance was still observed without significant changes to mean firing rates (Supplementary Fig. S2*F–H*).

An important reason behind the choice of studying the impact of HCN channels on synchrony propagation pertains to the critical roles that they play in regulating the operation mode, coincidence detection window, and spike initiation dynamics of neurons (Ratte et al., 2013; Das and Narayanan, 2014, 2015, 2017; Das et al., 2017). Specifically, owing to the slow negative feedback loop that HCN channels mediate (Mittal and Narayanan, 2024; Mishra and Narayanan, 2025), changes to their densities regulate neuronal ability to act as Class I integrators or Class II/III coincidence detectors (Ratte et al., 2013; Das and Narayanan, 2014, 2015, 2017; Das et al., 2017). We confirmed this with our models by demonstrating that in neurons with high HCN conductance, the STA displayed a narrow window associated with a negative lobe, indicating that these neurons preferentially responded to coincident inputs. In contrast, neurons with low-HCN density exhibited a broader and more slowly decaying STA, consistent with an integrator (Fig. 6*A*). As the ability of a neuron to perform coincidence detection constitutes an important requirement in synchrony detection (Softky and Koch, 1993; Softky, 1995; Konig et al., 1996; Singer et al., 1997; Engel et al., 2001; Hong et al., 2012; Ratte et al., 2013), we evaluated the impact of altering HCN-channel density on synchrony propagation across all assemblies of the feedforward network.

**Figure 6:**
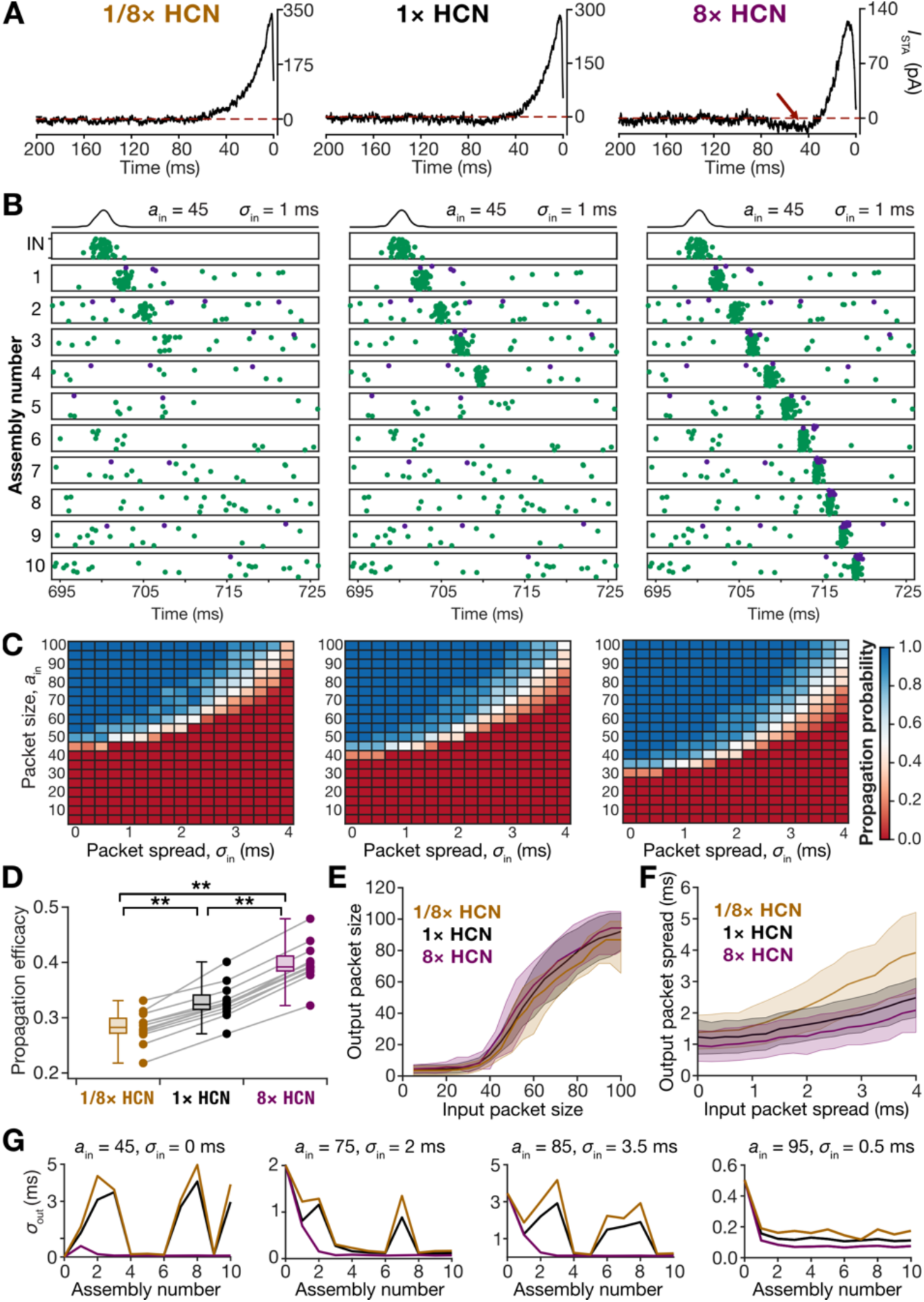
HCN channels enhanced success in synchrony propagation across homogeneous excitatory-inhibitory synfire chains. (A) Spike-triggered average (STA) of an excitatory neuron with low (1/8×), baseline (1×), and high (8×) HCN channel densities. Note the reduction in the peak currents and the emergence of a prominent negative lobe (depicted by an arrow in the 8× trace) with increase in density of HCN channels. (B) Increasing HCN conductance facilitated the propagation of spike volleys across the network for an example network receiving an input packet with *a*_*in*_ = 45 and *σ*_*in*_ = 1 ms. (C) Propagation probability matrices for an example network computed across 10 trials for each input packet size (*a*_*in*_) and spread (*σ*_*in*_), for low (left, 1/8×), baseline (middle, 1×), and high (right, 8×) HCN channel densities. Increase in HCN-channel density (left: 1/8×, 𝐸_*prop*_=0.27; middle: 1×, 𝐸_*prop*_=0.37; and right: 8×, 𝐸_*prop*_=0.43) shifted the stochastic separatrix, resulting in successful propagation of a larger set of input combinations. (D) Propagation efficacy 𝐸_*prop*_ plotted for 12 network initializations across three levels of HCN conductance. Lines connect the same network across the three HCN conductance values. **: *p*<0.005, Wilcoxon signed rank test. (E) Output packet size *a*_𝑜𝑢𝑡_ plotted for three different levels of HCN-channel conductances for different values of *a*_*in*_ (fixed *σ*_*in*_ = 1 ms). Thick lines are averages and shaded regions represent standard deviations. (F) Output packet spread *σ*_𝑜𝑢𝑡_ plotted for three different levels of HCN-channel conductances for different values of *σ*_*in*_ (fixed *a*_*in*_ = 50). Thick lines are averages and shaded regions represent standard deviations. (G) Output spread *σ*_𝑜𝑢𝑡_ plotted across all assemblies for a fixed input packet size with varying HCN-conductance densities (1/8×, 1×, and 8×).

To investigate the impact of HCN-channel conductance on synchrony propagation, we repeated our simulations with three different densities of HCN: a baseline condition with default HCN-channel density, a high HCN-channel condition where the density was increased eight-fold, and a low HCN-channel scenario in which the density was reduced to one eighth of the baseline. For each of these conditions, the simulations for 340 input combinations were repeated for 10 trials to assess trial-to-trial variability. These 10 trials of 340 input combinations were repeated for 12 independent network realizations to assess network-to-network variability. To build these independent homogeneous network realizations, different pairs of valid excitatory-inhibitory neurons (Figs. 2–3) were used to generate each assembly of each network.

Strikingly, we found that networks endowed with high values of neuronal HCN conductance showed enhanced propagation efficacy (Supplementary Fig. S3; Fig. 6*B–D*). We observed a progressive downward shift in the position of the stochastic separatrix with increasing HCN channel conductance, demonstrating that a larger number of input combinations successfully propagated in networks with high neuronal HCN conductance values (Fig. 6*C*). We quantified this using propagation efficacy of 12 different networks, each with 3 distinct scales of HCN-channel conductance. We found a significant increase in propagation efficacy across all twelve networks when neuronal HCN conductance values were elevated (Fig. 6*D*). This population-level consistency shows the generality of the effect, demonstrating that increasing HCN conductance reliably enhanced the ability of the network to propagate synchronous activity regardless of the specific intrinsic properties of the neurons chosen to define the homogeneous feedforward network.

Together, these results demonstrate that increasing HCN conductance enhances the ability of homogeneous feedforward networks to propagate synchronous activity.

### HCN-induced enhancement in synchrony propagation was mediated by the slow negative feedback dynamics of HCN channels

To understand why HCN conductance enhanced the propagation of synchrony in the feedforward chain, we first examined the input-output transformation performed by the first assembly of the network under the three HCN density conditions (Fig. 6*E–F*). When we compared the output volley size (*a*_𝑜𝑢𝑡_) for fixed input volley size (*a*_*in*_) and fixed input volley spread (*σ*_*in*_), we found that the mean *a*_𝑜𝑢𝑡_(across ten independent trials) remained largely comparable across the three HCN conditions (Fig. 6*E*). In striking contrast, when we examined the temporal spread of the output (*σ*_𝑜𝑢𝑡_) for fixed *a*_*in*_and *σ*_*in*_, we found that networks with higher HCN density showed reduced *σ*_𝑜𝑢𝑡_ compared to the low HCN and baseline conditions (Fig. 6*F*). This reduction indicates that the output activity of the first assembly was more tightly synchronized when HCN density is high. To investigate if this enhancement in temporal precision persisted beyond the first assembly, we tracked *σ*_𝑜𝑢𝑡_across all downstream assemblies for the same input combinations, but with different HCN-channel scaling (Fig. 6*G*). We observed this enhanced temporal locking observed with high HCN conductance to be preserved across the chain of assemblies and across input combinations, with assemblies in the high HCN networks consistently producing output packets with enhanced temporal coherence compared to than those in the low HCN networks (Fig. 6*G*). The enhanced temporal sharpening with high HCN-conductance provides a mechanistic explanation for the increased propagation efficacy observed in networks with elevated HCN conductance.

Spike initiation in neurons and the classes of excitability could be viewed to be emergent from interactions between fast and slow currents (Ratte et al., 2013; Mittal and Narayanan, 2024). Increase in HCN conductance results in an increase in the slow restorative current, thereby reducing the window within which output spikes can be generated (Ratte et al., 2013; Das and Narayanan, 2014, 2015, 2017; Mittal and Narayanan, 2024; Mishra and Narayanan, 2025). We hypothesized that this HCN-induced reduction in the window within which output spikes must be generated constituted the mechanistic basis for temporal sharpening in the output packet spread observed earlier (Fig. 6*F–G*). Within this framework, the slow kinetics of the restorative current constitutes an important attribute in defining this restricted window for spike generation, as the spike must be generated *before* the slow restorative current shuts down temporal summation (Ratte et al., 2013). Therefore, a nuanced test of our hypothesis on the mechanistic basis of how HCN-channels enhanced output synchrony would be to replace HCN channels with their faster counterparts without alteration to neuronal excitability (Das and Narayanan, 2014; Sinha and Narayanan, 2015). Specifically, if HCN channels were indeed enhancing output synchrony by reducing the window for spike generation through their slow kinetics, speeding up HCN kinetics without altering their gating properties (thereby retaining their restorative characteristic) should nullify the improvement in propagation efficacy achieved with high HCN-conductance.

In directly testing this, we next performed another set of simulations using a modified version of the HCN channel with faster activation kinetics. Specifically, we reduced the activation time constant (𝜏_*HCN*_) by a factor of twenty, which effectively removed the slow feedback dynamics that are normally mediated by HCN-channel activation and yielded a fast-HCN channel. This modification allowed us to isolate the specific contribution of the slow temporal dynamics to the propagation of synchronous activity. We then replaced the native slow HCN channels in all neurons in the network with these fast-HCN channels. The density of the fast-HCN channels was adjusted so that the neuron exhibited the same maximum impedance after replacement as it did with the native channels, thereby maintaining neuronal excitability as before. We repeated network propagations across all 340 input combinations, ten trials each, using baseline, eightfold, and one-eighth fast-HCN densities (Fig. 7*A*). Strikingly, when the native HCN channels were replaced by their faster counterparts, we no longer observed differences in propagation efficacy across all measured scales of HCN conductance (Fig. 7*A–B*; *cf*. Fig. 6*B–C* for the same network with same input combinations, but with slow HCN channels). The location of the separatrix remained comparable across the networks containing high, low, or baseline fast-HCN density. When we repeated these analyses across twelve independent network initializations, the results were consistent. We observed no significant change in propagation efficacy or location of the stochastic separatrix across different scales of HCN conductance (Fig. 7*C*).

**Figure 7:**
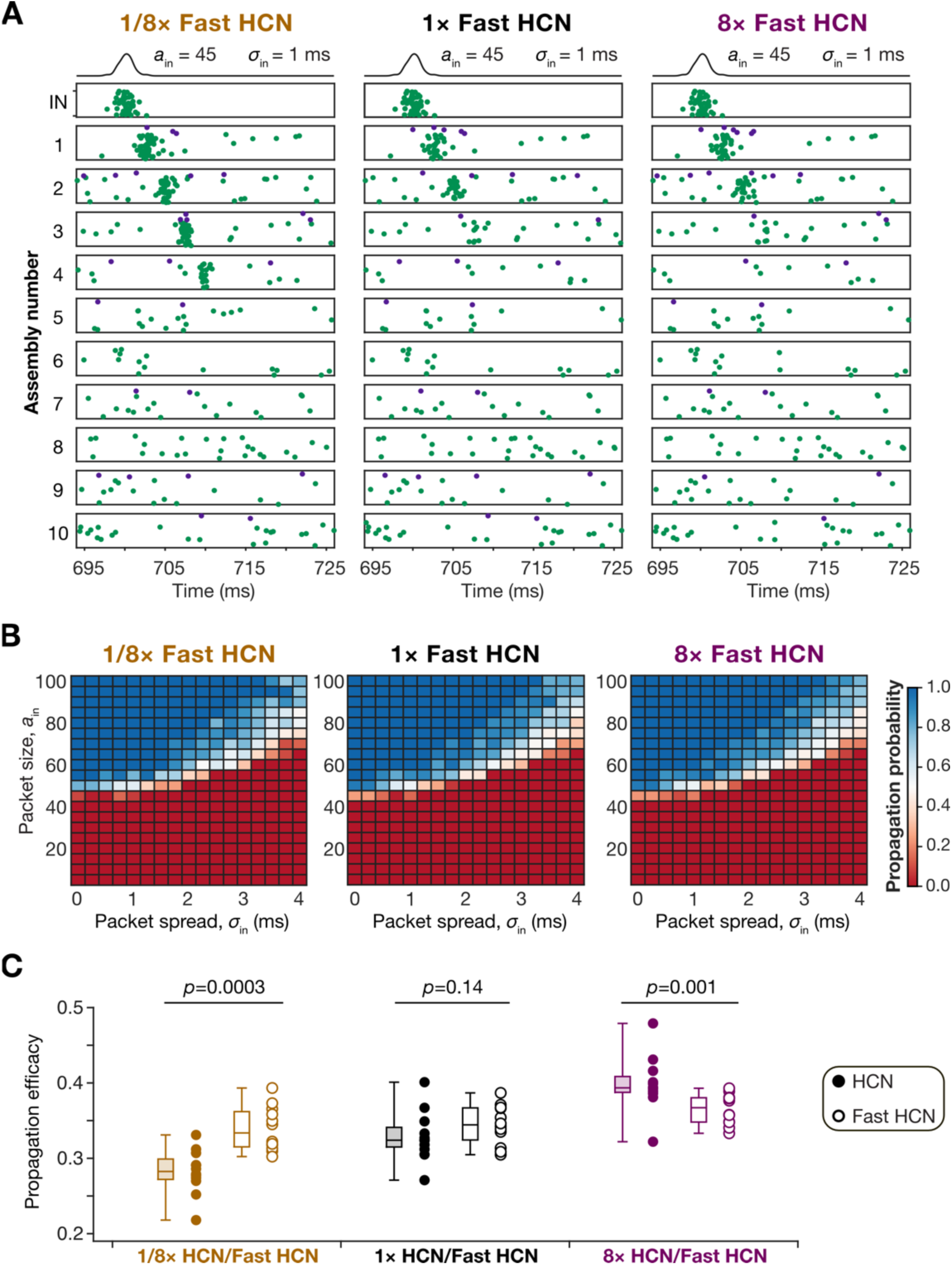
Fast-HCN channels did not alter synchrony propagation in synchrony propagation across homogeneous excitatory-inhibitory synfire chains. (A) Increasing conductance of HCN channels with fast kinetics did not significantly impact propagation of spike volleys across the network for an example network receiving an input packet with *a*_*in*_ = 45 and *σ*_*in*_ = 1 ms. Compare with Fig. 6B showing same network configuration with native slow HCN channels. (B) Propagation probability matrices for an example network computed across 10 trials for each input packet size (*a*_*in*_) and spread (*σ*_*in*_), for low (left, 1/8×), baseline (middle, 1×), and high (right, 8×) fast-HCN channel densities. Increase in density of fast HCN channels (left: 1/8×, 𝐸_*prop*_=0.33; middle: 1×, 𝐸_*prop*_=0.34; and right: 8×, 𝐸_*prop*_=0.34) did not markedly alter the stochastic separatrix. (C) Propagation efficacy 𝐸_*prop*_ plotted for 12 network initializations across three levels of HCN (same as Fig. 6D) or fast-HCN conductance. *p* values refer to outcomes of Wilcoxon signed rank test.

Together, these analyses provide direct mechanistic evidence in support of our hypothesis that the slow restorative nature of HCN channels mediates the restricted window for output spike generation, resulting in the lower output packet dispersion observed in the presence of high HCN-channel conductance (Fig. 6*E–G*). These results also demonstrate that the enhancement of synchrony propagation depends not simply on the presence of HCN conductance but on the intrinsic temporal characteristics of its activation kinetics, with the slow activation kinetics of the native HCN channels playing a crucial role in restricting the window for output spike generation.

### Intrinsic heterogeneities reduced network-to-network variability and HCN channels enhanced synchrony propagation in heterogeneous feedforward networks

Our analyses thus far have focused on homogeneous assemblies, where all 100 excitatory neurons and all 25 inhibitory neurons within each assembly were replicates of a single excitatory and a single inhibitory neuronal model, respectively. However, biological neural circuits are not composed of repeating homogeneous units. They are endowed with distinct forms of heterogeneities, which have been shown to play diverse and crucial physiological roles (Dahmen et al., 2026). Therefore, we assessed synchrony propagation under a more physiologically realistic scenario involving heterogeneous assemblies, specifically probing how intrinsic neuronal heterogeneity altered synchrony propagation across the network. Towards this, we constructed assemblies with graded degrees of heterogeneities, using the heterogeneous populations of excitatory and inhibitory neurons that we had generated (Figs. 2–4). These populations captured a wide range of intrinsic excitability and firing properties and provide the substrate for introducing controlled variability into the network.

We defined eight different degrees of heterogeneity ranging from the homogeneous degree 1, where a single excitatory and a single inhibitory neuron were used to build an assembly, to the completely heterogeneous degree 100, in which every neuron in the assembly was unique. Intermediate degrees of heterogeneity (2, 4, 8, 16, 32, and 64) were introduced by building each assembly with as many excitatory and inhibitory neurons. In general, for heterogeneity degree 𝐷_𝐻_(𝐷_𝐻_ ∈ {1,2,4,8,16,32,64,100}), there were 𝐷_𝐻_ number of unique excitatory models and 𝐷_𝐻_number of unique inhibitory models used to build every assembly of the network. For inhibitory neurons, when 𝐷_𝐻_ was greater than 25, all 25 inhibitory neurons in any assembly were unique. Thus, 𝐷_𝐻_ = 1 represents the homogeneous network studied thus far and 𝐷_𝐻_ = 100 represents a completely heterogeneous network. For each of the 8 degrees of heterogeneity, we simulated twelve independently initialized networks. For each network, we performed propagation across 10 trials for each of the 340 distinct (*a*_*in*_, *σ*_*in*_) combinations, computed propagation probability for each combination, and quantified propagation efficacy. To assess the impact of HCN channels on synchrony propagation in networks with different degrees of heterogeneity, we repeated this entire set of simulations for three scales of HCN-channel conductance (1/8×, 1×, and 8×), as before (Fig. 6).

We found pronounced network-to-network variability in propagation efficacy (Fig. 8, Fig. 9*A–C*), in the width of the stochastic separatrix (Fig. 8, Fig. 9*D–F*), and in the packet spread at the fixed point (Supplementary Fig. S4, Fig. 9*G–I*), even across different homogeneous networks (Fig. 8–9, Supplementary Fig. S4). Although there was no monotonic dependency of propagation efficacy on the degree of heterogeneities (Fig. 9*A–C*), we found some networks showing an increase in propagation efficacy with increasing degree of heterogeneity (*e.g*, Network #5 in Fig. 8) while others showed a reduction (*e.g.*, Network #8 in Fig. 8). Thus, there was network-to-network variability in how different networks reacted to introduction of graded heterogeneities as well (Figs. 8–9). We found that the mean propagation efficacy remained similar across all degrees of heterogeneity, showing that increasing heterogeneity did not systematically impair or enhance synchrony propagation (Fig. 9*A–B*). However, increasing degree of heterogeneity consistently reduced network-to-network variability in propagation efficacy (Fig. 9*C*). The mean or variance of the width of stochastic separatrix (Fig. 9*E–F*), the packet spread at the fixed point (Fig. 9*H–I*), and the packet size at the fixed point (Supplementary Fig. S5) across different networks did not manifest strong dependencies on the degree of heterogeneity.

**Figure 8:**
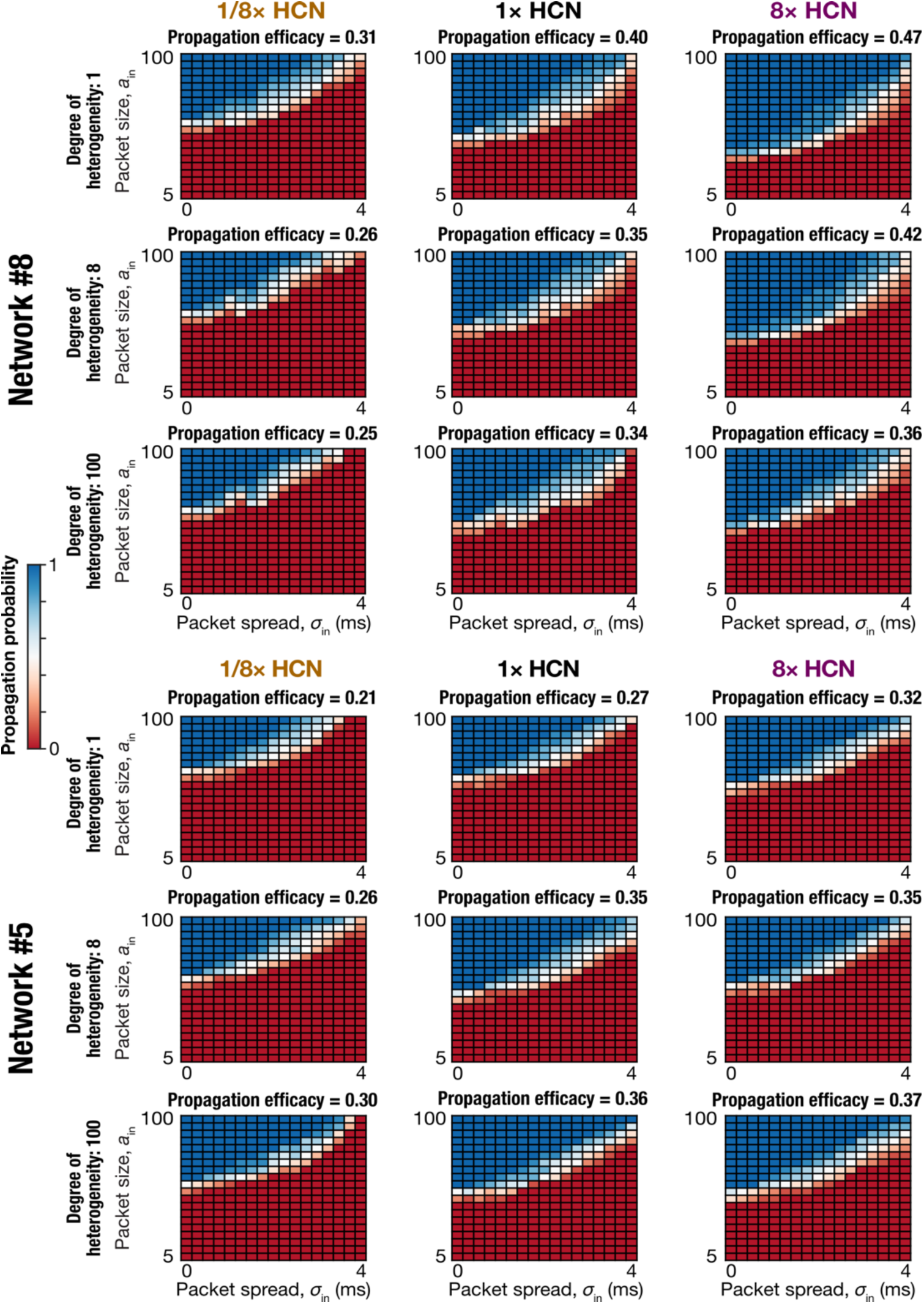
Network-to-network variability in synchrony propagation across graded levels of intrinsic heterogeneity and HCN-channel density values. Propagation probability matrices for two example networks (*Top*, Network #8; *Bottom*, Network #5) computed across 10 trials for each input packet size (*a*_*in*_) and spread (*σ*_*in*_), for low (left, 1/8×), baseline (middle, 1×), and high (right, 8×) HCN channel densities. Shown for each network are the matrices for three degrees of heterogeneity: 1 (*Top*), 8 (*Center*), and 100 (*Bottom*). It may be noted that propagation efficacy decreased for Network #8 but increased for Network #5, with increasing degree of heterogeneity. Increase in HCN-channel density consistently increased propagation efficacy across networks.

**Figure 9:**
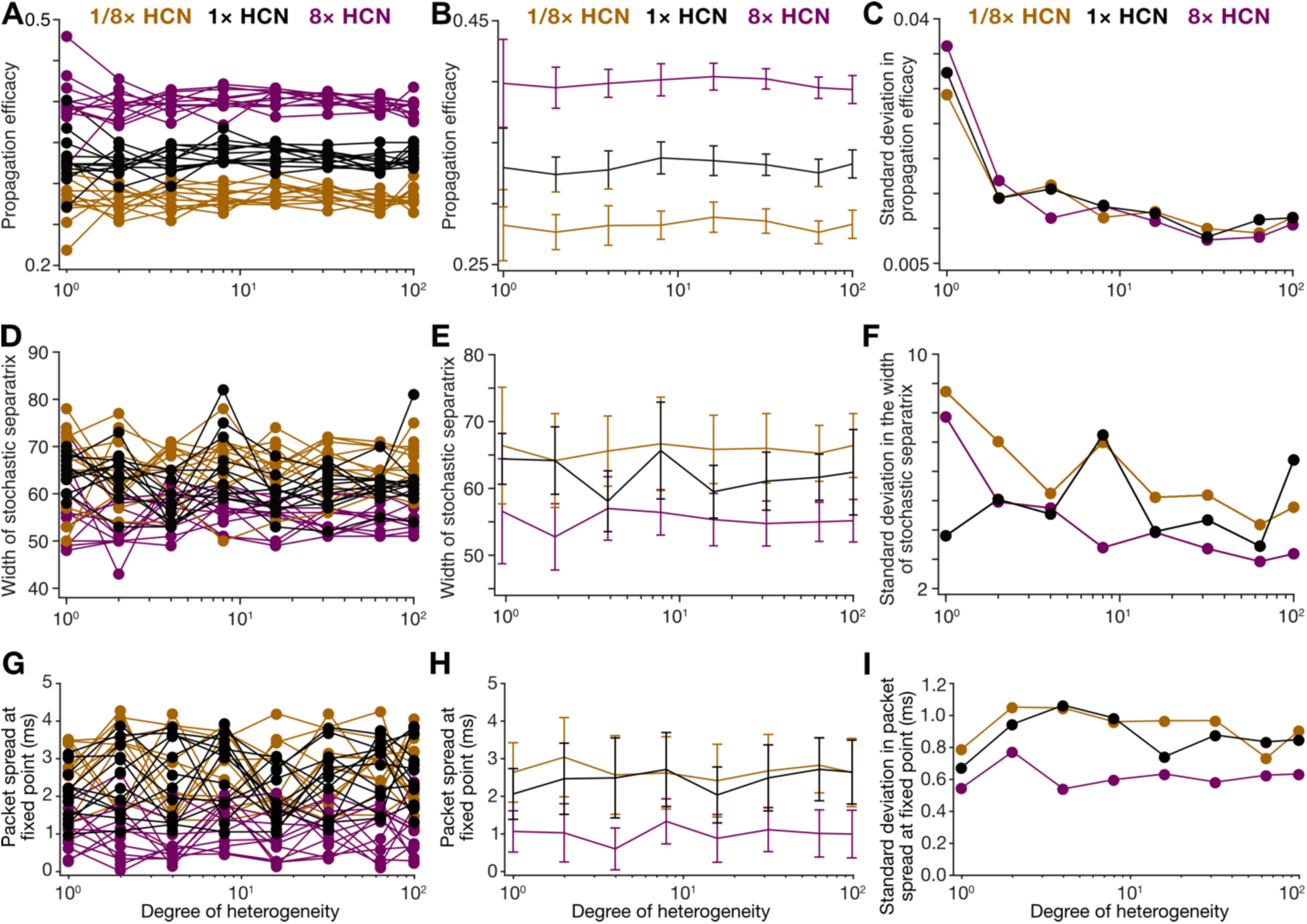
Increasing HCN-channel density enhanced propagation efficacy and reduced packet spread for successful propagations while enriched intrinsic heterogeneities reduced network-to-network variability in propagation efficacy. (A) Propagation efficacy plotted across 8 different degrees of heterogeneity for 12 independent network initializations and three levels of HCN-channel densities (1/8×, 1×, and 8×). (B) Mean propagation efficacy increased with HCN-channel densities but remained largely invariant to change in degree of heterogeneity. (C) Network-to-network variability in propagation efficacy decreased with increase in degree of heterogeneity but remained largely invariant to change in HCN-channel density. (D) Width of stochastic separatrix plotted across 8 different degrees of heterogeneity for 12 independent network initializations and three levels of HCN-channel densities (1/8×, 1×, and 8×). (E) Mean width of stochastic separatrix remained largely invariant to change in HCN-channel density or in degree of heterogeneity. (F) Network-to-network variability in the width of stochastic separatrix reduced with increase in degree of heterogeneity but remained largely invariant to change in HCN-channel density. (G) Packet spread at fixed point (convergence point for successful synchrony propagation through all ten assemblies) plotted across 8 different degrees of heterogeneity for 12 independent network initializations and three levels of HCN-channel densities (1/8×, 1×, and 8×). (H) Mean packet spread at fixed point reduced with increase in HCN-channel density but was invariant to change in degree of heterogeneity. (I) Network-to-network variability in the packet spread at fixed point was invariant to increase in degree of heterogeneity but reduced with increase in HCN-channel density.

Strikingly, across all degrees of heterogeneity and across all networks, increasing HCN channel density consistently increased propagation efficacy (Fig. 8, Fig. 9*A–B*), with minimal change to network-to-network variability (Fig. 9*C*). Although propagation efficacy consistently increased with increasing HCN-channel density across networks, there was network-to-network variability in the magnitude of such increase (Fig. 8). For any given density of HCN channels, the mean propagation efficacy across different networks was largely insensitive to increasing heterogeneity, whereas the variance across network realizations continued to decrease as heterogeneity increased (Fig. 9*A–C*). The mean or variance of the width of stochastic separatrix across different networks did not manifest strong dependencies on HCN-channel density either (Fig. 9*E–F*). However, both mean and variance of the packet spread at the fixed point reduced with increase in HCN-channel density (Fig. 9*H–I*), in a manner consistent with our previous observations on reduction in the window for spike generation with increase in HCN channels (Fig. 6). The mean or variance of packet size at the fixed point did not show strong dependencies on HCN-channel density (Supplementary Fig. S5).

Together, these results demonstrated pronounced network-to-network variability in propagation efficacy, in the width of the stochastic separatrix, and in the dependence on the degree of intrinsic heterogeneity. Introduction of intrinsic heterogeneity reduced network-to-network variability in propagation efficacy, without altering the average propagation efficacy across all degrees of heterogeneity. The most striking outcomes of our analyses were the consistent increase in propagation efficacy and the consistent reduction in packet spread at fixed point with increasing density of neuronal HCN channels, across different networks and across all degrees of heterogeneity.

## DISCUSSION

We systematically evaluated synchrony propagation in a large population of feedforward networks constructed from physiologically validated heterogeneous neuronal models spanning different degrees of intrinsic heterogeneity and distinct intrinsic properties. We generated heterogeneous valid model populations that faithfully captured experimentally observed variability in the intrinsic properties of cortical excitatory and inhibitory neurons. Our analyses revealed that signature neuronal functions could emerge from several non-random and non-unique combinations of functionally specialized molecular subsystems (ion channels and calcium buffering), together unveiling complexity and degeneracy at the cellular scale. We assembled these neuronal populations into feedforward networks with different degrees of heterogeneities, introducing background synaptic activity to ensure that neurons operated in a balanced high-conductance state and produced random spontaneous spiking activity. We studied synchrony propagation with a wide variety of input combinations, each assessed for multiple trials in different networks.

In homogeneous networks, we observed pronounced trial-to-trial variability in synchrony propagation for certain input combinations, giving rise to a stochastic separatrix, a zone of uncertainty that separated propagating and non-propagating input combinations. Our analyses demonstrated a strong dependence of propagation efficacy and the width of the stochastic separatrix on input packet spread, highlighting a strong dependence of trial-to-trial variability on input synchrony. Strikingly, increasing neuronal HCN conductance robustly enhanced the ability of the network to propagate synchronous activity. Mechanistically, we demonstrate that this enhancement was mediated by the slow restorative nature of HCN channels that restricts the temporal window for output spike generation, thereby reducing output packet dispersion when HCN-channel density was high. Thus, our analyses reveal that this enhancement depended not merely on the presence of HCN channels but relied critically on the slow activation kinetics of native HCN conductances, which played a central role in shaping the output timing window.

We introduced heterogeneities at different degrees in neuronal intrinsic properties of a population of feedforward networks and assessed their impact on synchrony propagation. With trial-to-trial variability still strong in heterogeneous networks, we found substantial network-to-network variability in propagation efficacy, width of the stochastic separatrix, and sensitivity to intrinsic heterogeneity. Importantly, introduction of intrinsic heterogeneities reduced network-to-network variability in propagation efficacy, without altering the mean propagation efficacy.

Extending from our analyses with homogeneous networks, we found a consistent increase in propagation efficacy with increasing neuronal HCN channel density across different networks and all degrees of intrinsic heterogeneity. Together, our analyses demonstrate the ability of disparate combinations of heterogeneous neurons with distinct molecular composition to yield similar synchrony propagation efficacy across feedforward networks. Such network-scale degeneracy in synchrony propagation, coupled with cellular-scale degeneracy in achieving characteristic neuronal properties, point to degeneracy as a fundamental organizing principle in the emergence of neural-circuit function.

### HCN channels are active regulators of synchrony propagation

The prominent and most consistent outcome of our analyses is that increasing neuronal HCN channel density increased the propagation efficacy across all networks that we tested, irrespective of the degree of heterogeneity. Our analyses also provide biophysical mechanistic insights into why this was the case by demonstrating a critical role for the slow kinetics of the negative feedback loop mediated by HCN channels. In essence, in the presence of HCN channels, the synchronous inputs need to elicit spikes *before* the slow negative feedback loop limits further spike generation. With increasing HCN channel density, the strength of the negative feedback increases with the spike-generation window becoming narrower. The narrow spike generation window implies that the *output packet spread* of an assembly reduces with increasing HCN channel density (Fig. 6*F–G*, Supplementary Fig. S4, Fig. 9*H–I*). This characteristic feature of HCN channels manifests as the strong negative lobe in the spike triggered average, which also reduces the integration/coincidence detection window in the presence of HCN channels, together yielding coherent output spikes (Das and Narayanan, 2014, 2015; Sinha and Narayanan, 2015; Das and Narayanan, 2017; Das et al., 2017; Mishra and Narayanan, 2025). The inability of fast HCN channels to yield similar enhancement in synchrony propagation (Fig. 7) provides direct support for a role of the slow negative feedback motif in mediating HCN-induced increase in synchrony propagation efficacy.

Our results indicate that, as a direct consequence of their unique biophysical characteristics that allowed them to enhance output synchrony, HCN channels regulated the position of the stochastic separatrix that separates propagating from non-propagating input packets. Specifically, increasing HCN-channel density shifted the separatrix, expanding the range of input combinations that successfully propagated through the network, whereas reducing HCN narrowed this range. These observations are important from two well-established signatures associated with cortical neurons. First, the dendrites of cortical pyramidal neurons are highly enriched with HCN channels (Williams and Stuart, 2000; Berger et al., 2001; Kole et al., 2006; Kalmbach et al., 2013). As the first layer of neuronal integration for inputs, active dendritic structures can convey synchrony information to the cell body through dendritic spikes or plateau potentials and their timing. Thus, the high density of HCN channels in the dendrites offers an effective strategy to improve synchrony propagation efficacy.

Second, HCN channels are strongly regulated by activity-dependent plasticity and by several neuromodulatory systems (including serotonin, dopamine, noradrenaline, and acetylcholine) which influence arousal, attention, and behavioral state (McCormick and Pape, 1990; Pape, 1996; Fan et al., 2005; Rosenkranz and Johnston, 2006; Narayanan and Johnston, 2007; Rosenkranz and Johnston, 2007; Narayanan and Johnston, 2008; Dembrow et al., 2010; Narayanan et al., 2010; Dembrow and Johnston, 2014; Shah, 2014; Santello and Nevian, 2015; Labarrera et al., 2018; Shah, 2018; Mishra and Narayanan, 2025). Depending on receptor expression and the specific downstream signaling cascades, different activity-dependent plasticity mechanisms and disparate neuromodulators can have region-specific effects on HCN conductance. Through such regulation, plasticity or neuromodulation could shift the stochastic separatrix, selectively enhancing or reducing the propagation of synchronous activity, in a behavioral state-dependent manner. These routes provide physiologically plausible mechanisms by which the cortex can dynamically route information according to behavioral context by altering a specific neuronal conductance that is highly expressed in cortical structures.

Together, HCN-dependent control of the stochastic separatrix represents a flexible mechanism for regulating information flow by dynamically altering the priority for synchronous activity, for efficient routing and gating of afferent signals across cortical microcircuits. This perspective emphasizes that the impact of HCN extends beyond single-cell characteristics. By modulating the network-scale boundary between propagating and non-propagating inputs, HCN channels provide a tunable lever for controlling the flow of information in cortical circuits. Neuromodulatory and activity-dependent regulation of HCN channels can thus implement a context-dependent filter, dynamically shaping which inputs are transmitted across layers and which are suppressed, ultimately influencing perception, attention, and cognitive processing.

### Stochastic separatrix of synchrony propagation in high-conductance networks points to functional roles for background activity

Our study shows the manifestation of a stochastic separatrix, whereby an identical input packet in the exact same network propagates in some trials but not in others. These observations provide evidence that the propagation of synchronous activity in high-conductance networks depends not just on the network or the characteristics of the input packet, but also on the specific ongoing background activity. The state of the network at the specific moment that an input packet arrives could either facilitate or impede propagation, effectively setting an additional gate on whether input synchrony is transmitted or not. Specifically, background synaptic activity introduces stochastic fluctuations in membrane potential and in neuron gain (through the specific set of receptor conductances that contribute to the gain). The specific background activity patterns that are coincident with the afferent input packet bring about temporally targeted changes to the membrane potential and neuronal gain, thus contributing directly to whether spike generation could occur for that input packet. As a result, even identical input packets could propagate in some trials but fail in others, depending on the temporal alignment of the membrane potential and the gain aligned with the afferent input packet. These observations highlight that propagation is not solely determined by neuronal intrinsic properties and the characteristics of the input packet but is also highly sensitive to the temporal context provided by ongoing background synaptic activity.

These observations point to potentially important functional roles for background synaptic activity in cortical circuits. Different forms of synaptic fluctuations, which could take the form of correlated patterns of inputs onto neurons, oscillatory fluctuations at different frequencies, or temporally structured synaptic barrages, could shift the position or widen the range of the stochastic separatrix. Importantly, the potential conversion of non-propagating input patterns to propagating ones in such noisy conditions could potentially give rise to stochastic resonance in synchrony propagation, whereby an optimal level of noise actively enhances the propagation of borderline input combinations. These conclusions lend further credence to observations across biological systems that noise is not merely a source of variability or interference but serves as a dynamic regulator of information flow (Bulsara et al., 1991; Longtin, 1993; Wiesenfeld and Moss, 1995; Gluckman et al., 1996; Simonotto et al., 1997; Russell et al., 1999; Stacey and Durand, 2001; Mori and Kai, 2002; Schneidman et al., 2003; Moss et al., 2004; McDonnell and Abbott, 2009; Levchenko and Nemenman, 2014; Selimkhanov et al., 2014; Mittal and Narayanan, 2022).

Cortical circuits could thus exploit this principle by modulating the amplitude, temporal structure, or correlation of background synaptic activity, effectively creating a tunable gate for synchrony propagation. Such mechanisms could allow circuits to selectively amplify certain inputs while suppressing others, depending on behavioral state, attentional focus, or neuromodulatory context. Furthermore, this perspective suggests a rich interplay between deterministic input properties and stochastic network dynamics. Background activity sets the operating point of neurons and determines how close they are to firing threshold at the exact moment an incoming packet arrives. In this sense, the network’s noise environment could prime circuits for either reliable propagation or selective filtering, introducing an additional layer of control over information transmission. Future studies could systematically explore how different noise statistics influence the stochastic separatrix, investigate if specific patterns of background activity could enhance signal reliability, and whether these effects could be harnessed by the brain to regulate the routing of information in a structured and context-dependent manner. The implications for such analyses to synchrony propagation in brain regions where there are multiple bands of behaviorally relevant oscillatory activity would provide further insights about this noise-mediated gate to synchrony propagation (Buzsaki, 2006, 2010; Sinha and Narayanan, 2015; Mittal and Narayanan, 2022; Sinha and Narayanan, 2022).

### Degeneracy as a foundational organizing principle to achieve flexible and robust function in heterogeneous cortical neural circuits

Intrinsic heterogeneity is a fundamental feature of biological systems across all scales. Apart from other functions that intrinsic heterogeneities play, they form the substrate for degeneracy, allowing multiple combinations of parameters to yield similar functional outcomes. In our study, the heterogeneous populations of excitatory and inhibitory neurons that we generated using the MPMOSS algorithm spanned the full parametric range for each conductance and exhibited weak pairwise correlations across parameters (Figs. 2–4). These results demonstrate cellular-scale degeneracy in both neuronal subtypes, showing that functionally valid neurons emerge from widely different sets of intrinsic conductances, highlighting the degeneracy inherent in these cell types.

If any single parameter defining excitatory or inhibitory neurons were individually chosen, their value across the population would span a large range (Fig. 2*B*, Fig. 3*B*), underlining parametric heterogeneity therein. However, the important observation here is that heterogeneity in *any* individual parameter, when analyzed *in isolation* doesn’t provide insights about valid model populations. This is because a large proportion of model samples that were generated from the identical distribution for the specific parameter were deemed invalid by the MPMOSS algorithm. Only 2.6% of excitatory neurons and 6.2% of inhibitory neurons, generated from the exact same distribution of individual parametric values, were deemed valid. Thus, from the perspective of that single parameter, the extent of heterogeneity in the valid and the invalid model subpopulations would be identical. Thus, *heterogeneities in individual parameters* do not offer insights into how cellular function was achieved. It is instead the *specific combinations* of *all parameters* that yield models that satisfy characteristic cellular physiology. Importantly, it is not a single combination or combinations that are clustered around a single combination that yield valid models, but combinations endowed with wide distributions of all underlying parameters that matched characteristic cellular physiology (Figs. 2–4). This ability of disparate combinations of functionally independent subsystems (*e.g.,* different ion channels and buffers) that yield integrated function of the collective system is referred to as degeneracy (Edelman and Gally, 2001). Together, the specific combinations of subsystems that yield characteristic cellular physiology are neither random (as inferred from the large percentage of invalid models) nor unique (follows from degeneracy), placing single neurons as complex systems built of disparate functionally specialized subsystems (Seenivasan and Narayanan, 2022; Albantakis et al., 2024; Kumari and Narayanan, 2024; Mittal and Narayanan, 2024). An important observation here is that studying heterogeneity in any single component is insufficient to provide deep insights about the system, instead emphasizing the need to focus on the global structure of the parametric space.

At the network scale, our work demonstrates that propagation efficacy of networks depends on the characteristics of the afferent input packet, the intrinsic properties of neurons in the network, the constituent set of neurons that form the network (yielding network-to-network variability), the degree of intrinsic heterogeneity in assemblies, and the background activity coincident with the input packet (mediating trial-to-trial variability). Thus, if synchrony propagation is the collective function that the network is to implement, there are disparate combinations of functionally specialized subsystems that yield similar functional outcomes. This network-scale degeneracy that is built with cellular subsystems (neurons) that were built from molecular subsystems through cellular-scale degeneracy points to a cascade of degeneracy (Albantakis et al., 2024) in the emergence of network function. Thus, the disparate forms of heterogeneities and specific interactions among them form the substrate that mediates flexible circuit function with degeneracy as the foundational organizing principle in heterogeneous cortical circuits.

Our results show that the average synchrony propagation efficacy is largely invariant to the degree of within-subtype intrinsic heterogeneity. This indicates that cortical circuits can reliably preserve synchronous transmission despite substantial heterogeneities at the single-cell level. Heterogeneous networks therefore maintain a balance between flexibility and stability, consistently producing functional outputs while accommodating a wide range of intrinsic cellular properties. These findings suggest that the heterogeneity observed in cortical circuits may serve as a source of robustness, allowing networks to maintain reliable function despite perturbations or fluctuations in individual neurons and supporting stable computation in a dynamically changing environment. Overall, heterogeneities in molecular and cellular components together ensure that cortical networks can operate robustly through disparate specific combinations of underlying molecular and cellular components.

### Limitations and future directions

While our study provides insights into how HCN channel and intrinsic heterogeneity regulate synchrony propagation in cortical microcircuits, rigorously accounting for stochasticity and variability in propagation, there are important limitations to the approach. First, owing to the computational complexity of network simulations, our network model is built of single-compartmental models of cortical excitatory and inhibitory neurons. The tremendous computational capabilities of cortical neuronal dendrites of both excitatory and inhibitory neurons are well-established (Johnston et al., 2003; Johnston and Narayanan, 2008; Major et al., 2013; Tzilivaki et al., 2019; Larkum, 2022). With differential densities of channels across the dendritic arbor (Williams and Stuart, 2000; Berger et al., 2001; Kole et al., 2006; Harnett et al., 2013; Kalmbach et al., 2013), each contributing to the spike-initiation dynamics differentially (Prescott et al., 2006; Prescott et al., 2008a; Prescott et al., 2008b; Ratte et al., 2013; Das and Narayanan, 2014, 2015, 2017; Das et al., 2017), it is essential that the dendritic arbor and its active components are explicitly accounted into future analyses. In addition, as cortical pyramidal neurons receive inputs from different brain regions onto different parts of their dendritic arbor, it is critical that these distinct inputs and spatiotemporal patterns activity therein are explicitly accounted for.

Second, in our study we focused specifically on manipulating HCN channels to examine their impact on synchrony propagation. However, neurons express a wide variety of voltage- and calcium-gated ion channels that contribute to excitability, subthreshold resonance, and spike timing (Hutcheon and Yarom, 2000; Narayanan and Johnston, 2012; Ratte et al., 2013; Das et al., 2017; Rathour and Narayanan, 2019; Mittal and Narayanan, 2024). Future studies could explore how modulation of other conductances, alone or in combination with HCN, influences the propagation of synchronous activity, potentially revealing additional mechanisms for tuning network-level information flow. Of particular interest here might also be interactions between feedforward inhibition and HCN channels, either of which could act as slow negative feedback loop motifs, in restricting the spike generation window and enhancing synchrony propagation. Third, our manipulation of HCN channels was uniform across neurons, whereas in the brain HCN expression is heterogeneous and subject to spatial and neuromodulatory gradients. Investigating how localized or neuron-specific modulation of HCN affects network-level propagation could reveal mechanisms for selective routing of information or attention-dependent gating. Finally, while our study provides testable predictions on the role of HCN channels, background synaptic activity, and intrinsic heterogeneities, experimental validation *in vivo* or *in vitro* will be essential. Targeted modulation of HCN channels combined with measurements of synchronous activity under controlled input conditions could confirm the predicted shifts in the stochastic separatrix and the enhancement of temporal precision. Similarly, manipulating intrinsic heterogeneity or background synaptic statistics could probe the robustness and reproducibility of synchronous propagation in real cortical circuits. With reference to intrinsic heterogeneities, for instance, certain pathological conditions have been shown to result in loss of heterogeneities (Rich et al., 2022). This provides a substrate for assessing the impact of pathological conditions in general and heterogeneities in particular, towards regulating synchronous propagation across the cortical network.

In summary, while our model captures key biophysical and network principles underlying synchrony propagation, extending it to incorporate dendritic spikes and plateau potentials driven by such synchrony, patterned background activity, and spatially heterogeneous neuromodulation will be essential for fully understanding how cortical circuits dynamically regulate information flow under physiological conditions. Future studies could also explore the impact of stochasticity in gating of ion channels and heterogeneities in synaptic properties on synchrony propagation as well, while the network *learns* to mediate synchrony propagation through conjunctive changes in intrinsic and synaptic properties.

## ACKNOWLEDGMENTS

The authors thank Dr. Arvind Kumar, Dr. Sufyan Ashhad, and members of the cellular neurophysiology laboratory for helpful discussions and for comments on a draft of this manuscript. This work was supported by the Ministry of Education (R. N. & S. S.).

## DATA AVAILABILITY STATEMENT

The data underlying this study are available as part of the article. The original source code used for the simulations in this article will be made publicly available upon article acceptance.

## Author Contributions

S. and R. N. designed experiments; S. S. performed experiments; S. S. analyzed data; S. S. and R. N. wrote the paper.

