## Supplementary Figures S1-S5 for "HCN channels enhance synchrony propagation in heterogeneous synfire chains"

### Supplementary Material

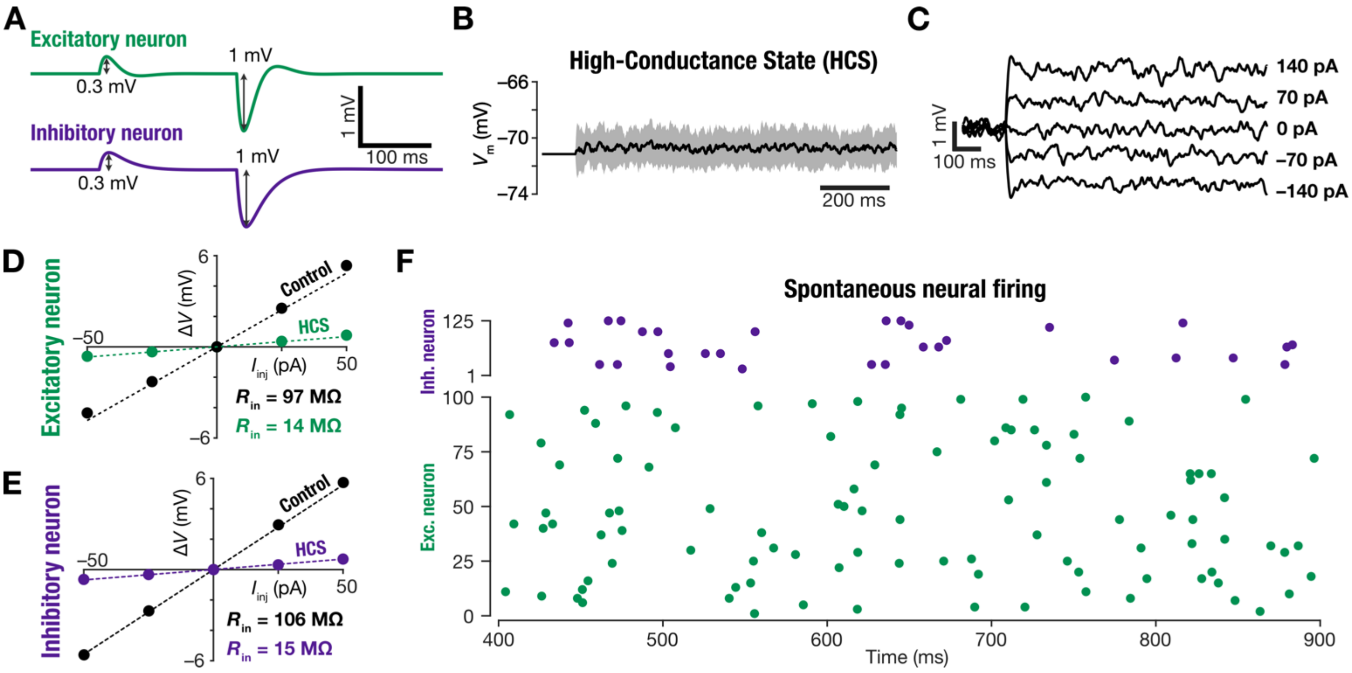

**Supplementary Figure S1: High-conductance states regulate neuronal intrinsic properties.** (A) Excitatory and inhibitory synaptic weights were tuned for each neuron to produce an excitatory postsynaptic potential (EPSP) of 0.3 mV and an inhibitory postsynaptic potential (IPSP) of 1 mV. (B) Each neuron received 5,000 randomized background synaptic inputs, 80% of which were excitatory (2 Hz) and 20% inhibitory (2.2 Hz). The black trace shows the mean membrane potential across 10 trials in the presence of background synaptic activity. The shaded area represents the standard deviation. (C) Voltage-current ($V - I$) relationship of an excitatory neuron under step current injections ($-140$ to $140$ pA) in the presence of background synaptic inputs. Each trace reflects the average across 10 trials. (D–E) Input resistance of an excitatory (D) and an inhibitory (E) neuron reduced in the presence of background synaptic inputs, indicative of a high-conductance state induced by synaptic receptor activation. (F) Random spontaneous spiking activity in the excitatory and inhibitory neuronal populations following the addition of randomly timed, high-strength synaptic inputs to each neuron. Shown are the spontaneous spike timings over a 500 ms period after the initial 400 ms stabilization period.

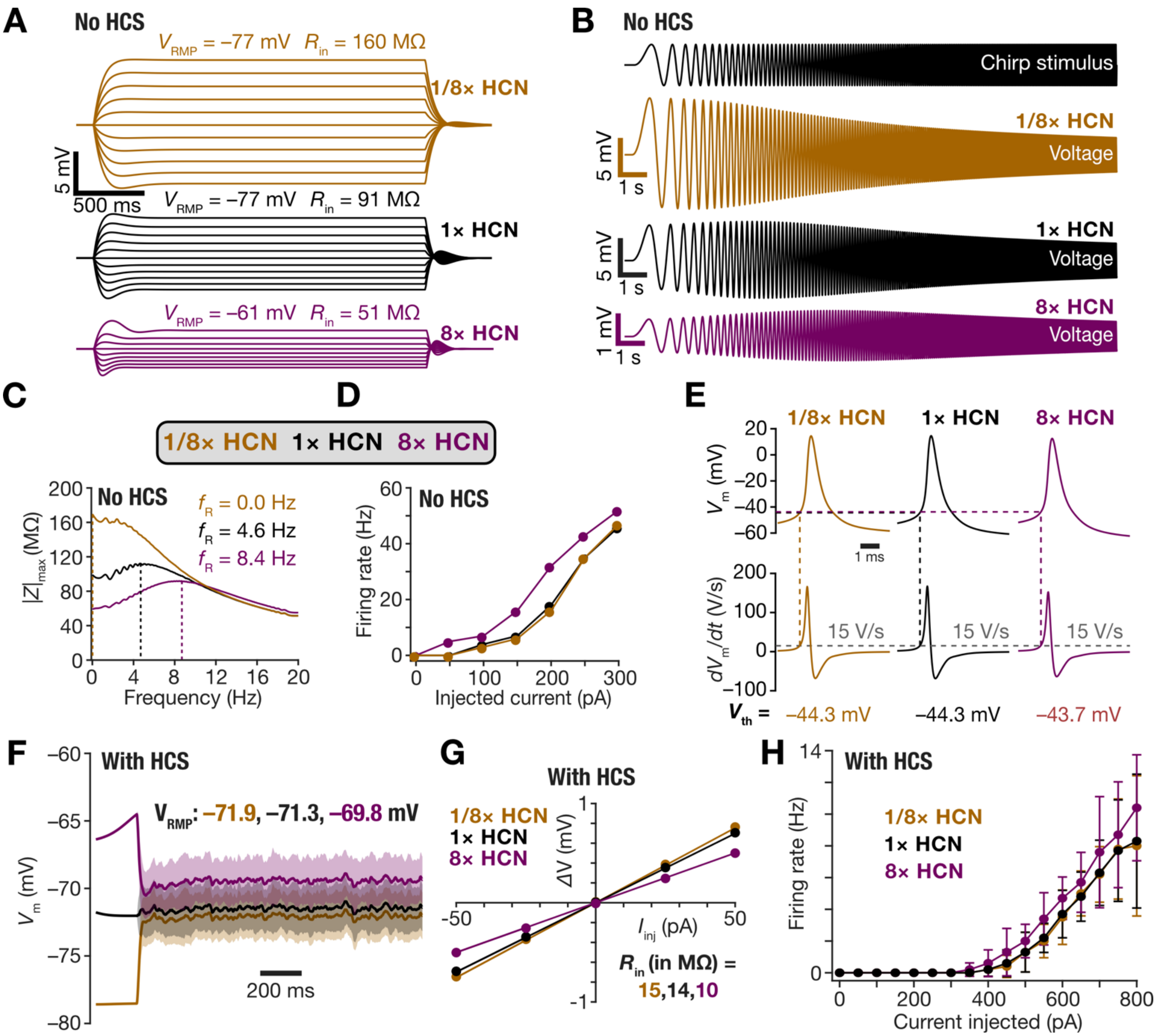

**Supplementary Figure S2: Regulation of neuronal intrinsic properties by HCN channels in the presence *vs.* absence of high-conductance state.** (A–E) In the absence of external activity and the consequent high-conductance state (HCS), increasing HCN channel density depolarized the resting membrane potential and reduced the input resistance (A), decreased the maximum impedance amplitude, and increased the resonance frequency (B–C), increased the firing rate (D), but did not significantly alter action potential threshold characteristics (E). (F–H) In the presence of high-conductance state, increasing HCN-channel density minimally depolarized the average resting membrane potential across 10 trials (F), reduced the input resistance (G), and did not significantly increase the firing rate across 10 trials (H). The minimum current required to elicit action potentials increased in the presence of high-conductance state (panel D *vs*. panel H), across all densities of HCN channels. It may be also noted that while RMP depolarized by ~16 mV in the absence of high-conductance state (panel A), the average RMP shifted by ~1 mV in the presence of high-conductance states (panel F). All measurements were obtained at the respective RMP, after the RMP settled to a relative steady state.

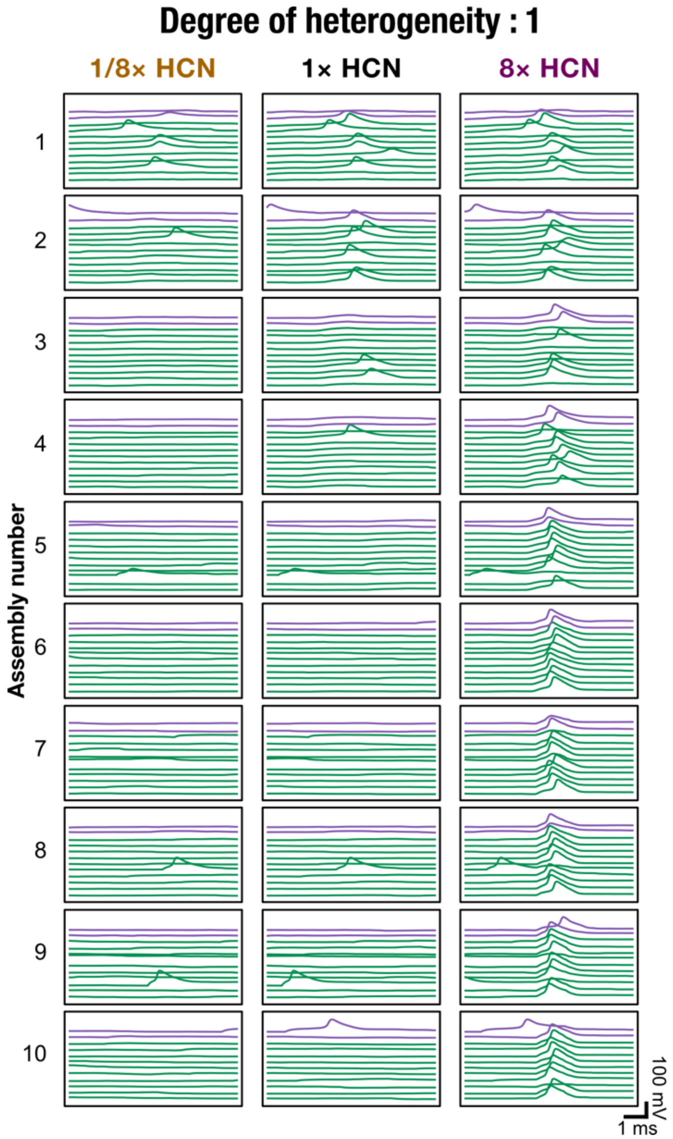

**Supplementary Figure S3: Dependence of synchrony propagation on graded levels of intrinsic heterogeneity and HCN-channel density values.** Representative voltage traces from 10% of all excitatory (10 green traces in each assembly) and inhibitory neurons (2 purple trace in each assembly) for two degrees of intrinsic heterogeneity (1 and 100) and three levels of HCN-channel densities (1/8×, 1×, and 8×). Strong synchrony propagation might be noted for 8× HCN traces irrespective of the degree of heterogeneity.

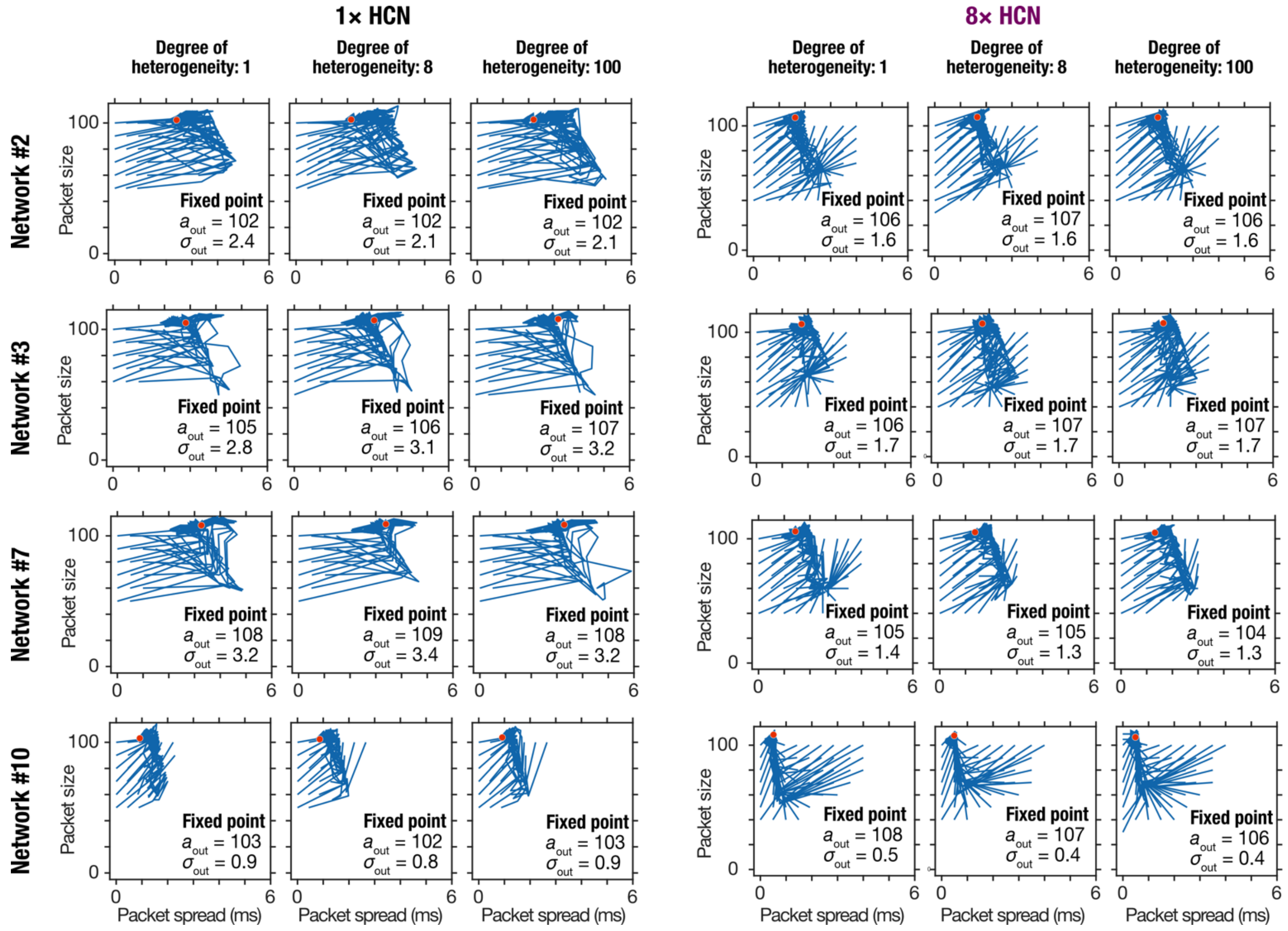

**Supplementary Figure S4: Dependence of fixed point for successful synchrony propagation through all assemblies on graded levels of intrinsic heterogeneity and HCN-channel density values.** Representative phase plane trajectories for all successful synchrony propagation across all 10 assemblies, shown for 4 networks, three degrees of intrinsic heterogeneity (1, 8, and 100), and two levels of HCN-channel densities (1× and 8×). The orange circles represent the fixed points where the trajectories converge. The value of output packet size and spread at the respective fixed points are provided as insets in each phase plane plot. Across networks, it may be noted that while the packet spread at the fixed point reduced with increase in HCN-channel density, it was invariant to change in degree of heterogeneities. There was no discernible dependence of packet size at the fixed point as functions of HCN-channel density or degree of heterogeneities.

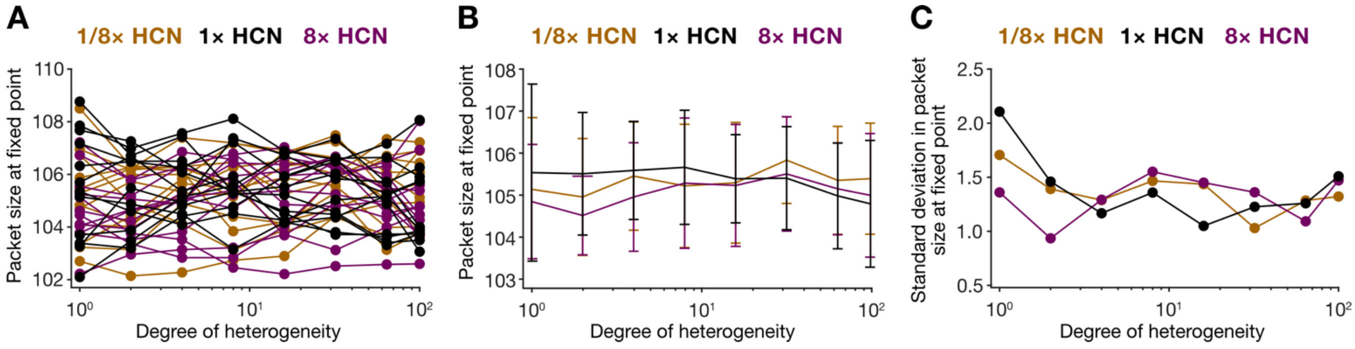

**Supplementary Figure S5: Packet size at fixed point for successful synchrony propagation through all assemblies was largely invariant to changes in HCN-channel density or in degree of intrinsic heterogeneities.** (A) Packet size at fixed point (convergence point for successful synchrony propagation through all ten assemblies) plotted across 8 different levels of heterogeneity for 12 independent network initializations and three levels of HCN-channel densities (1/8×, 1×, and 8×). (B) Mean packet size at fixed point was invariant to increase in HCN-channel density or in degree of heterogeneity. (C) Network-to-network variability in the packet size at fixed point was invariant to increase in degree of heterogeneity or in HCN-channel density.
